# DEVELOPMENTAL AND EVOLUTIONARY COMPARATIVE ANALYSIS OF A REGULATORY LANDSCAPE IN MAMMALS AND BIRDS

**DOI:** 10.1101/2021.08.12.456039

**Authors:** Aurélie Hintermann, Isabel Guerreiro, Christopher Chase Bolt, Lucille Lopez-Delisle, Sandra Gitto, Denis Duboule, Leonardo Beccari

## Abstract

Modifications in gene regulation during development are considered to be a driving force in the evolution of organisms. Part of these changes involve rapidly evolving *cis*-regulatory elements (CREs), which interact with their target genes through higher-order 3D chromatin structures. However, how such 3D architectures and variations in CREs contribute to transcriptional evolvability remains elusive. During vertebrate evolution, *Hox* genes were redeployed in different organs in a class-specific manner, while maintaining the same basic function in organizing the primary body axis. Since a large part of the relevant enhancers are located in a conserved regulatory landscape, this gene cluster represents an interesting paradigm to study the emergence of regulatory innovations. Here, we analysed *Hoxd* gene regulation in both murine vibrissae and chicken feather primordia, two mammalian- and avian-specific skin appendages which express different subsets of *Hoxd* genes, and compared their regulatory modalities with the regulations at work during the elongation of the posterior trunk, a mechanism highly conserved in amniotes. We show that in the former two structures, distinct subsets of *Hoxd* genes are contacted by different lineage-specific enhancers, likely as a result of using an ancestral chromatin topology as an evolutionary playground, whereas the regulations implemented in the mouse and chicken embryonic trunk partially rely on conserved CREs. Nevertheless, a high proportion of these non-coding sequences active in the trunk appear to have functionally diverged between the two species, suggesting that transcriptional robustness is maintained despite a considerable divergence in CREs’ sequence, an observation supported by a genome-wide comparative approach.

## INTRODUCTION

Changes in the spatial and temporal regulation of genes critical for developmental processes have greatly contributed to the evolution of animal morphologies (see for example (Carroll, 2008; Rebeiz et al., 2015). The expression of such genes is generally modulated by combinations of *cis-*regulatory elements (CREs), short DNA sequences enriched for transcription factor binding sites, which tend to evolve more rapidly than the genes they control (Edwards et al., 2013; Long et al., 2016; Spitz and Furlong, 2012). CREs can be spread over large distances around the gene(s) they regulate, which gives to the spatial organization of chromatin a particular importance. Indeed, the existence of such ‘regulatory landscapes’ (Spitz et al., 2003) implies that enhancer-promoter interactions involve spatial proximity, which is generally thought to occur through large chromatin loops (Rao et al., 2014).

The extent of regulatory landscapes often correspond to Topologically Associating Domains (TADs) (Andrey et al., 2013; Harmston et al., 2017), which were defined as chromatin domains where the probabilities of interactions, as measured by chromosome conformation capture (Belton et al., 2012), are higher than when compared to the neighbouring regions (Dixon et al., 2012; Nora et al., 2012; Sexton et al., 2012). The distribution of TADs tends to be conserved in various vertebrate species, a phenomenon likely associated with regulatory constraints exerted by the complex and pleiotropic regulations found around many vertebrate developmental genes. Such domains, where many interactions might occur, were proposed to form particular chromatin niches, rich in various DNA-binding proteins and where the evolutionary emergence of novel regulatory sequences could be favoured due to the presence of both a pre-existing scaffold and appropriate co-factors (see (Darbellay and Duboule, 2016a). While genomic rearrangements altering the structure of TADs were reported to impact proper gene regulation during development, sometimes leading to genetic syndromes (e.g. (Bolt et al., 2021a; Bompadre and Andrey, 2019; Lupianez et al., 2015), in most cases, these rearrangements also affected enhancer-promoter spatial relationships, which makes a causal assessment difficult.

Genome-wide analyses carried out in different species, comparing DNA sequence conservation, TAD organization, or some epigenetic modifications, have started to reveal some of the molecular mechanisms underlying both the evolution of gene regulation and, by extension, the modification of species-specific traits. However, integrated functional approaches are required to better understand how changes in chromatin architecture and modifications in CREs may influence the evolvability and the robustness of gene transcription. To try to address such questions, we have used the *HoxD* locus as a paradigm of pleiotropic regulation of a multi-gene family, with a strong impact both on the control of important developmental steps, and on the emergence of evolutionary novelties. The *HoxD* cluster contains a series of genes in *cis,* which encode transcription factors that are expressed in different combinations across several embryonic structures. The cluster is flanked by two large TADs, each one matching a gene desert highly enriched in CREs (Darbellay and Duboule, 2016a). These two TADs are separated by a boundary region which is localized within the cluster, close to its centromeric extremity (Andrey et al., 2013; Rodriguez-Carballo et al., 2017). The telomeric TAD comprises a large number of enhancers controlling transcription of various *Hoxd* genes in most of the tissues where HOX proteins operate (Andrey et al., 2013; Darbellay et al., 2019; Delpretti et al., 2013; Guerreiro et al., 2016; Schep et al., 2016). In contrast, the centromeric TAD contains enhancers specific for terminal structures such as digits or external genitals (Amândio et al., 2020; Montavon et al., 2011).

Amongst the many tissues where *Hoxd* genes exert a function are embryonic primordia of different skin derivatives, including those of mammary glands, vibrissae and pelage hairs of mammals as well as those of chicken feathers (Kanzler et al., 1997, 1994; Reid and Gaunt, 2002; Reynolds et al., 1995; Schep et al., 2016)(Fig. 1). Despite the fact that these ectodermal structures share common developmental mechanisms and gene regulatory networks such as the *Wnt,* EDA and BMP signaling pathways (Biggs and Mikkola, 2014; Dhouailly et al., 2019; Di-Poï and Milinkovitch, 2016; Pantalacci et al., 2008; Wu et al., 2004), each type of primordium expresses different subsets of contiguous *Hoxd* genes. Since the specific topological organization of the *HoxD* locus can be observed in fishes (Acemel et al., 2016) (see Fig. S1A), it likely predates the emergence of hairs and feathers during amniote evolution (Dhouailly, 2009; Wu et al., 2004). For this reason, these organs represent valuable models to understand how CRE evolution within a pre-existing chromatin architecture could lead to the implementation of new gene-specific expression patterns. In contrast, the general organization of *Hoxd* gene expression along the main embryonic body axis is overall maintained across vertebrate species, in spite of the low sequence conservation observed, raising the question as to how evolving *Hox* clusters could maintain globally similar transcriptional outputs.

**Figure 1.**
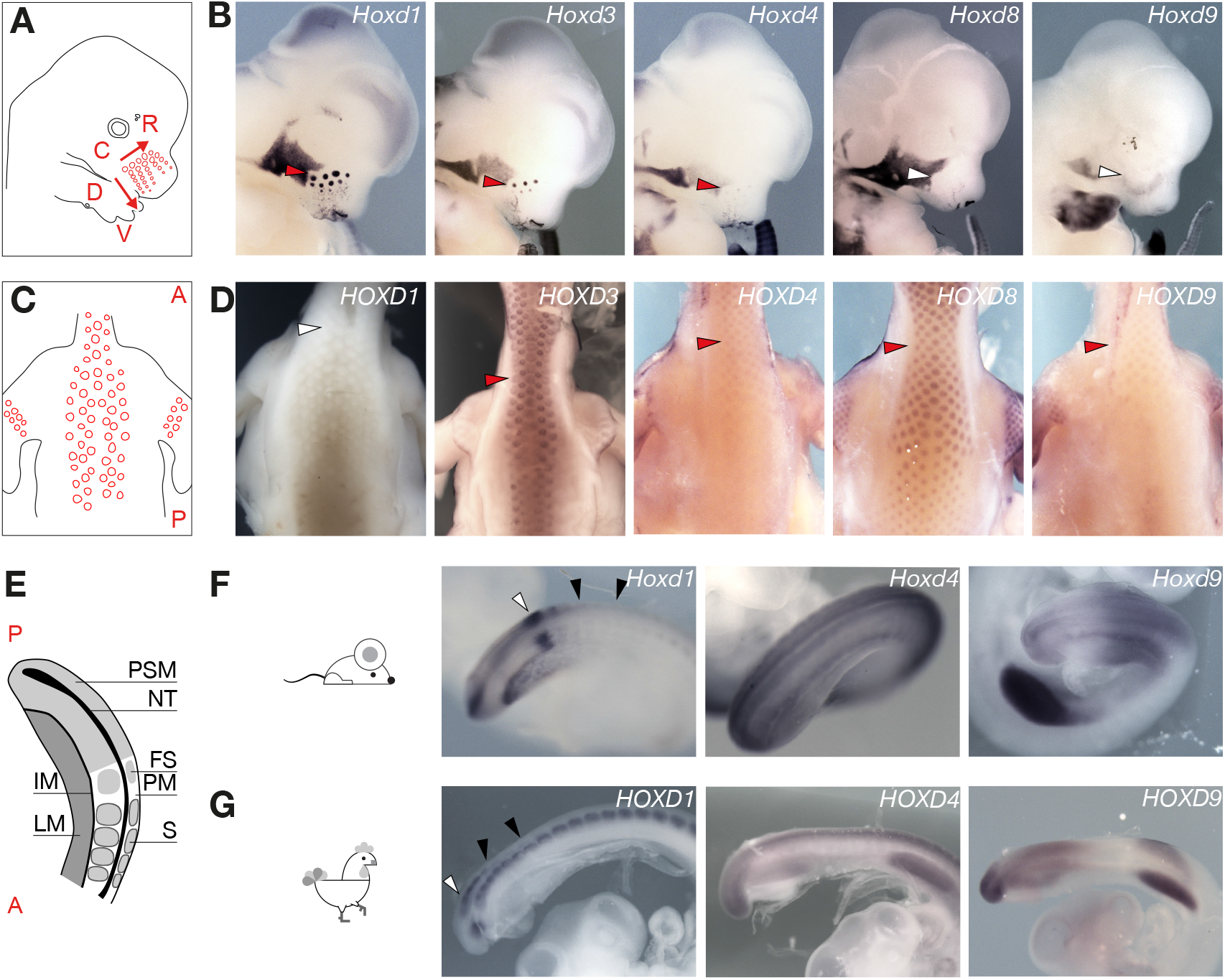
Transcription of *Hoxd* genes in mammalian and avian skin primordia. **A.** Schematic illustration of a E12.5 mouse embryo with VPs (Vibrissae Primordia) represented as red circles. The dorso-ventral (D-V) and the caudo-rostral (C-R) axes are shown as red arrows representing the timing of appearance of the vibrissae primordia. **B.** WISH on E12.5 faces. The arrowheads point to the most caudo-dorsal placode, which is the largest one and the first to appear. Staining intensity progressively decreases from *Hoxd1* to *Hoxd8.* **C.** Schematic illustration of the back of a HH35 chicken embryo with feather placodes as red circles. A, anterior; P, posterior. **D.** WISH on HH35 chicken showing the skin of the upper back. The arrowheads point to the neck at the level of the shoulders. *HOXD1* is not expressed, whereas *HOXD3* and *HOXD8* signal intensities are stronger than for *HOXD4* and *HOXD9.* **E.** Schematic representation of a E9.5 or HH20 posterior trunk and tailbud. A, anterior; P, posterior; PSM, presomitic mesoderm; NT, neural tube; IM, intermediate mesoderm; FS, forming somites; PM, paraxial mesoderm; LM, lateral mesoderm; S, somites. **F.** WISH on E9.5 mouse posterior trunk and tail bud. *Hoxd1* transcription pattern is different from that of *Hoxd4* and *Hoxd9.* **G.** WISH on HH20 chicken posterior trunk and tail bud. *HOXD1* transcription pattern is different from *HOXD4* and *HOXD9.* The white arrowheads in panels **F** and **G** point to the *Hoxd1* stripe in the PSM. The black arrowheads show the difference between mouse (**F**) and chicken (**G**) in the *Hoxd1* mRNAs persistence in formed somites.

In this study, we addressed these issues by taking a comparative look at a well-defined regulatory landscape, positioned telomeric to the *HoxD* gene cluster, in mammals and in birds. These two syntenic regions contain CREs involved either in the same regulatory tasks, or controlling regulatory aspects specific to each taxon. In the latter context, we characterized *Hoxd* gene regulation in the mouse-specific vibrissae primordia (VPs) and in the chicken-specific feather primordia (FPs). We show that different subsets of contiguous *Hoxd* genes are expressed in the mouse VP mesoderm and in the chicken FP ectoderm, indicating an independent co-option of *Hoxd* paralogs in these structures. While the transcription of these genes in VPs and FPs relies on lineage-specific CREs located within different segments of the regulatory landscape, these CREs exploit a largely conserved chromatin 3D architecture.

Our comparative analysis of the regulation at work during trunk extension also revealed that *Hoxd1* displays an expression pattern markedly different from that of its neighboring paralogs, both in mouse and chick, and that this *Hoxd1*-specific expression is driven by an evolutionary conserved CRE located in a region predominantly interacting with *Hoxd1* in both species. However, the cis-regulatory code at work in the latter regulation differs between the two species, illustrating the possibility that inter-species conservation of both gene transcriptional specificity and chromatin architectures are compatible with a high plasticity of the *cis*-regulatory sequences involved. The extension of this observation genome-wide suggests that conserved CRE sequences more frequently diverge in their regulatory activities than they conserve their functionality.

## RESULTS

### Different *Hoxd* gene expression in mammalian and avian skin appendages

We initially analyzed the expression profiles of *Hoxd* genes in mouse VPs and chicken FPs (Fig. 1). Skin appendage development starts with the interaction of the skin ectoderm with its underlying mesoderm, resulting in the thickening of the epithelium into a placode and in the condensation of dermal cells into a papilla (Mikkola, 2007; Sawyer and Knapp, 2003; Widelitz and Chuong, 1999). VP development starts at early embryonic day 12 (E12), forming a stereotyped array of eight columns and five rows of VPs (Wrenn and Wessells, 1984)(Fig. 1A). The epithelial placode subsequently invaginates and engulfs the dermal papilla forming the follicle primordium (Fig. S1). Likewise, in birds, ectoderm-mesoderm interactions result in the evagination of feather buds around embryonic day 7 (stage HH30-32 (Hamburger and Hamilton, 1951; Michon et al., 2007) (Fig. S2).

Whole-mount *in situ* hybridization (WISH) revealed strong *Hoxd1* expression in the dermal papillae of the VPs (Fig. 1B; Fig. S1). Furthermore, a comparative analysis of VPs from single E12.5 embryos revealed that *Hoxd1* transcripts accumulate slightly after ectodermal placode formation, marked by the expression of *Shh* (Chiang et al., 1999) (Fig. S1). *Hoxd3* and *Hoxd4* are also expressed in VPs, yet at much lower levels than *Hoxd1* (Fig. 1B, Fig. S1). Faint levels of *Hoxd8* mRNAs were sporadically detected in the VPs, while *Hoxd9* and more 5’-located *Hoxd* mRNAs were never observed in these structures (Fig. 1B, Fig. S1). *In situ* hybridization on cryostat sections confirmed that *Hoxd1* transcripts accumulate in the dermal mesenchyme of VPs, but not in the ectoderm-derived portion of the follicle (Fig. S1). Therefore, *Hoxd1*-*Hoxd4* are transcriptionally active in the dermal papillae of developing VPs, with *Hoxd1* being the main paralog expressed in these structures.

WISH experiments in HH35 chicken embryos revealed expression of *HOXD3, HOXD4, HOXD8* and *HOXD9* in the FPs, while *HOXD1* and *HOXD11* mRNAs were not detected (Fig. 1C, D, Fig S2). Cryostat sections indicated that *HOXD* gene expression in chicken FPs is localized in the follicle ectoderm (Fig. S2), in contrast with murine VPs. Different *Hoxd* genes thus operate in dermal papillae of mouse VPs and in ectodermal placodes of chicken FPs, likely reflecting the implementation of different regulatory mechanisms in the two lineages.

### *Hoxd* gene expression in the mouse and chicken embryonic body axis

*Hox* genes are expressed in the paraxial and lateral mesoderm of the main embryonic axis, as well as in the neural tube, and display a progressively more restricted expression domain depending on their position within the cluster (Gaunt et al., 1988). Comparative WISH analysis of mouse and chicken embryos revealed that *Hoxd1* expression patterns in the trunk were markedly similar between the two species, yet largely divergent from that of the 5’-located neighbor *Hoxd* paralogs (Fig. 1E-G). While *Hoxd4* was strongly and uniformly transcribed in the mouse paraxial and lateral mesoderm as well as in the neural tube, *Hoxd1* displayed a biphasic expression pattern. Transcripts accumulated in the embryonic tailbud, yet they rapidly disappeared from the presomitic mesoderm and remained undetected in lateral mesoderm progenitors or in the neural tube (Fig. 1F). Instead, *Hoxd1* was specifically re-activated during somite condensation, coupling the transcription of this gene to the segmentation clock (Dale and Pourquié, 2000; Zákány et al., 2001). Similar to the mouse pattern, the chicken *HOXD1* expression also differed from its neighbouring paralogs, even though its transcripts remained detectable in the formed somites all along the main body axis (Fig. 1G). These observations suggested that *Hoxd1* transcription in developing somites may depend on a specific set of CREs, different from CREs controlling the neighboring paralogs, and that this locus-specific transcription was evolutionarily conserved across amniotes.

### An evolutionary conserved topology at the *HoxD* locus

In order to see whether these differences and similarities in regulatory specificities could be associated to variations in the global chromatin architecture at this locus, we performed Capture Hi-C (CHi-C) experiments using dissected mouse and chicken posterior trunks (Fig. 2A, B). Even though the global size of TADs at this locus is approximately two times smaller in the chicken genome, the *HOXD* region closely resembled its mouse counterpart, with the T-DOM being further organized into two sub-TADs in both species (Fig. 2, sub-TAD1 and sub-TAD2) (Rodríguez-Carballo et al., 2020; Yakushiji-Kaminatsui et al., 2018). Of note, the presence and relative distribution of conserved non-coding elements (CNEs) were maintained within this similar chromatin architecture (Fig. 2, Fig. S3).

**Figure 2.**
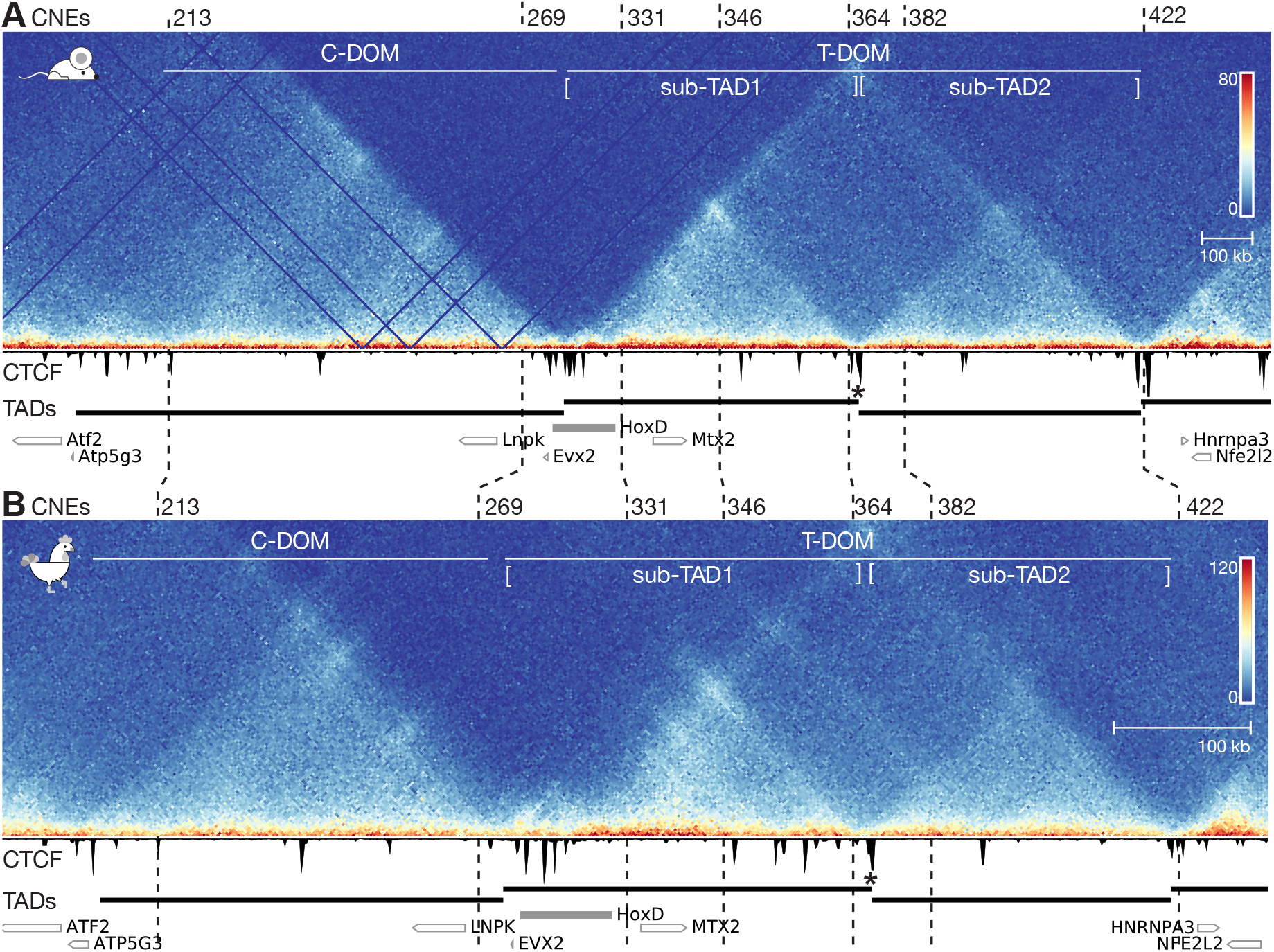
Capture Hi-C-seq at the mouse and chicken *HoxD* loci. **A, B.** Capture Hi-C-seq heatmaps using dissected posterior trunk cells. Below each heatmap, a CTCF ChIP-seq track is shown, produced from the same material. The similarities in the structural organization of the mouse and the chicken loci are underlined either by the positions of syntenic CNEs at key positions (numbered vertical dashed lines), the domains produced by TAD calling (black bars below) and the presence of a sub-TAD boundary (asterisk) within T-DOM. Genes are represented by empty rectangles. In panel **A,** E9.5 mouse posterior trunk cells were used and the CHi-C heatmap was mapped on mm10 with a bin size of 5 kb (chr2:73800000-75800000). In panel **B,** HH20 chicken posterior trunk cells were used and the CHi-C heatmap mapped on galGal6 with a bin size of 2.5kb (chr7:15790000-16700000) and on an inverted x-axis. The positions of the two TADs (T-DOM and C-DOM) and of both sub-TADs are shown on top. The scales on the X axes were adjusted to comparable sizes for ease of comparison, yet the chicken locus is more compacted (see the scale bars, bottom right).

We corroborated these observations by assessing the interactions established by the *Hoxd1, Hoxd4* and *Hoxd9* promoters in mouse and chicken embryonic posterior trunk cells, using 4C-seq (Fig. S3). As an indication for non-tissue-specific interactions, we used embryonic brain cells where *Hox* genes are not transcribed. In both species, the three viewpoints established frequent contacts with the T-DOM while interactions with the C-DOM were virtually absent. In addition, all viewpoints contacted T-DOM sub-TAD1 with a higher probability than T-DOM sub-TAD2, showing specific and non-overlapping regions where interactions were enriched. *Hoxd1* contacted the first half of sub-TAD1 (Figs. 3A and S3, region D1), more intensively than *Hoxd4* or *Hoxd9* (Fig. S3). In turn, *Hoxd4* contacts within the 3’ half of sub-TAD1 (Figs. 3A and S3, region D4), were increased up to the sub-TAD boundary (Fig. S3). Finally, *Hoxd9* interactions were mostly limited to the sub-TAD boundary region (Figs. 3A and S3, region D9), where they were higher than for *Hoxd1* and *Hoxd4.* This distribution of interacting regions within the 3’ TAD were found in both brain and posterior trunk cells, yet with an overall lower contact frequency in brain cells, reflecting the transcription-independent default conformation of the region, in agreement with previous reports (Andrey et al., 2013; Dekker and Heard, 2015; Noordermeer et al., 2011).

**Figure 3.**
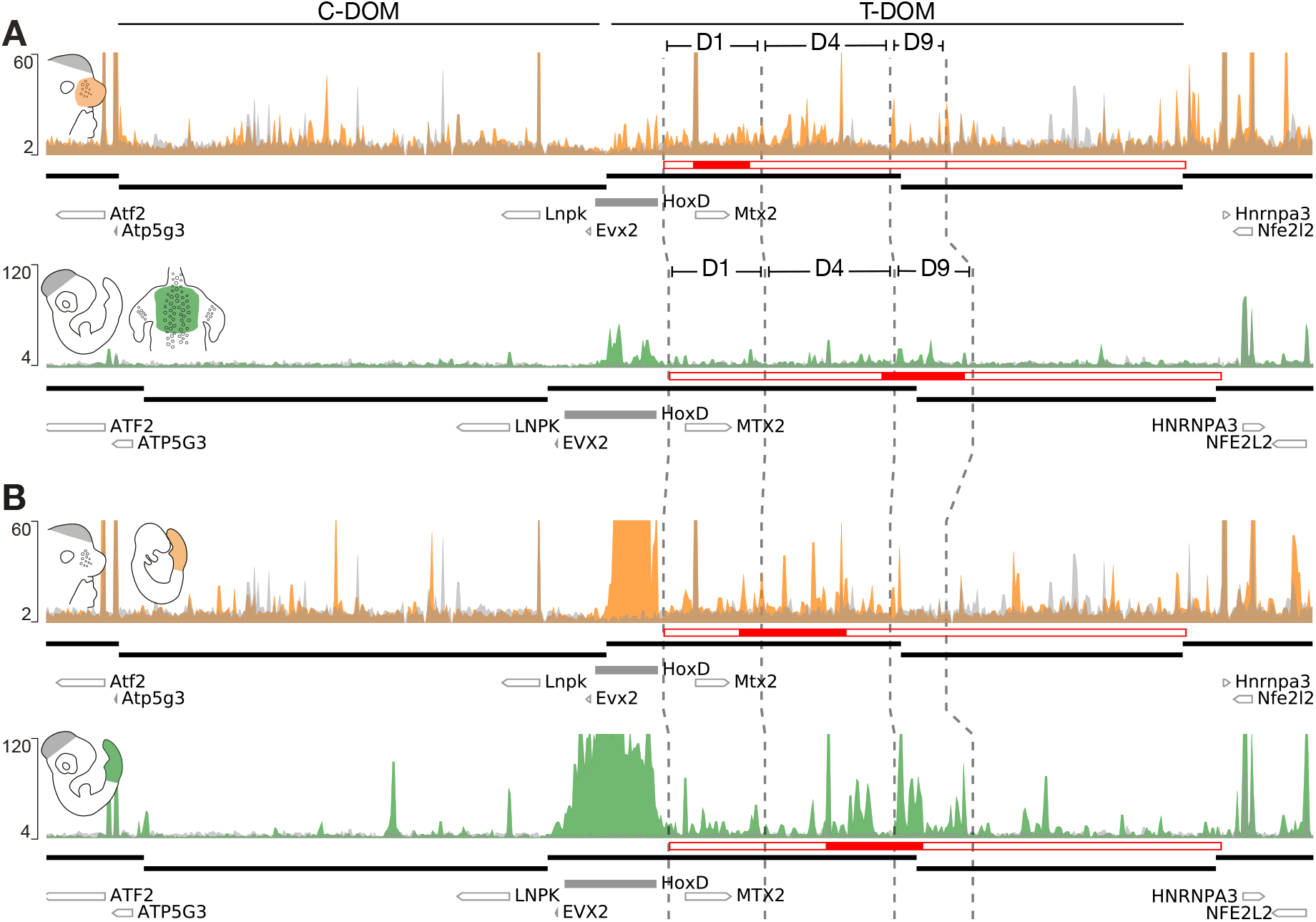
Tissue- and gene-specific interactions of dense H3K27ac regions. **A.** On top is a ChIP-seq profile of H3K27ac using mouse E12.5 VPs (orange) superimposed over the E12.5 forebrain cells (grey) (mm10, chr2:73800000-75800000). Below is a H3K27ac profile produced from dissected chicken HH35 dorsal skin (green), with HH18 brain cells (grey)(galGal6, chr7:15790000-16700000, inverted x-axis). The highest interacting regions are depicted as D1, D4 and D9 (see also Figure S3B). The positions of conserved CNEs were used to delimit these regions in both species (vertical dashed lines) and the extents of TADs are shown below (thick black lines). The T-DOM (empty red rectangle) is split in overlapping genomic windows and the densest window (Most Acetylated Region; MAR) is shown as a filled red rectangle. In the mouse VP, the MAR is within the D1 DNA segment, whereas the chicken MAR overlaps with the D4 and D9 regions. **B.** H3K27ac ChIP-seq using mouse E9.5 posterior trunk cells (orange) superimposed over forebrain cells (grey)(mm10, chr2:73800000-75800000). Below is shown the H3K27ac profile obtained from the same sample but dissected from a HH18 chicken embryo (green) with brain cells (grey) as a control (galGal6, chr7:15790000-16700000, inverted x-axis). While the mouse posterior trunk Most Acetylated Region (MAR, red rectangle) coincides with the D4 segment, the chicken MAR counterpart also covers D4 and a small part of D9.

Chromatin architectures partly rely upon the presence of the zinc-finger DNA binding protein CTCF, which together with the cohesin complex can induce the formation and stabilization of large loops. (Hansen et al., 2017; Nanni et al., 2020; Pugacheva et al., 2020). The ChIP-seq analysis of the CTCF profiles revealed that both at the mouse and chicken *HoxD* loci, the telomeric end of the D1 region maps close to an occupied CTCF site (Fig. S3, blue arrowhead). Also, a cluster of CTCF binding sites delimits the boundary between the mouse and the chicken sub-TAD1 and sub-TAD2 (Rodriguez-Carballo et al., 2017; Rodríguez-Carballo et al., 2020; Yakushiji-Kaminatsui et al., 2018) thus correlating with the limit between the *Hoxd4* and *Hoxd9* interacting D4 and D9 regions in both species (Fig. 2; Fig. S3, blue arrowheads in panel B).

Overall, these results show that *Hoxd1, Hoxd4* and *Hoxd9* preferentially interact with different portions of the T-DOM and that these segments are somehow determined by the distribution of CTCF binding sites in T-DOM subTAD-1. These distinct interactions patterns are constitutive and conserved between mouse and chicken. From these results, we concluded that the evolutionarily conserved contact topology of *Hoxd* genes with T-DOM may contribute to the implementation of their specific regulatory strategies in mouse and chicken embryonic structures.

### Tissue-specific regulatory sequences are located in regions of preferential interactions

To evaluate whether the promoter-specific partition of contacts with T-DOM is associated with the presence of differential regulatory activities, we set up to identify the CREs controlling *Hoxd* gene expression in the VPs, FPs and the embryonic trunk of both mouse and chicken embryos. We carried out ChIP-seq analyses using an antibody against H3K27ac, a histone modification indicative of an active chromatin state (Creyghton et al., 2010; Rada-Iglesias et al., 2011) and annotated tissue-specific regions with increased density of H3K27ac enrichment referred to as MARs (Most Acetylated Regions, Fig. 3). MARs were identified as genomic regions accounting for the highest H3K27ac enrichment after subtraction of the corresponding signal in brain cells (see material and methods).

In the mouse VPs, the MAR mapped within the D1 region (Fig. 3A, top, red-filled rectangle), in agreement with the stronger expression of *Hoxd1* compared to its neighbouring paralogs (Fig. 1B; Fig. S1). Instead, in chicken dorsal skin samples where *HOXD1* is not active (Fig. 1D), the MAR largely overlapped the D4 and the D9 interacting regions (Fig. 3A, bottom, red-filled rectangle). Therefore, VPs and FPs display different distributions of H3K27ac enrichment across T-DOM, in agreement with the various sibsets of *Hoxd* gene expressed in these tissues. In contrast, the MARs of both mouse and chicken posterior trunk samples largely overlapped the D4 region (Fig. 3B, red-filled rectangles) as expected from their similar expression patterns in the embryonic trunk of both species, with the expression domain of *Hoxd4* being more extended than that of *Hoxd9* (Fig. 1F). In summary, the different enrichments of H3K27ac marks across T-DOM were located within the genomic segments that preferentially interacted with the genes expressed in each tissue.

### *Hoxd1* regulation in mutant regulatory landscapes

To evaluate the functional importance of the D1 interacting region, we assessed *Hoxd1* transcription in different mouse lines carrying mutations affecting this regulatory domain. We first analyzed *Hoxd1* expression in the VPs of mouse embryos carrying various deletions of T-DOM (Fig. 4A, red lines). In the *HoxD^Del(attP-TpSB2)lac^* allele, the D1 region is removed (hereafter referred to as *Del(attP-SB2).* Since *Del(attP-SB2)* homozygous embryos are not viable due to the deletion of the essential *Mtx2* gene (Andrey et al., 2013), we crossed this line with the mouse line *HoxD^Del(1-13)d9lac^* (Spitz et al., 2001), hereafter referred to as *Del(1-13).* In the latter line, the entire *HoxD* cluster is deleted, such as to be able to see the effect of the *HoxDD^el(attP-SB2)lac^* deletion upon *Hoxd* gene regulation in *cis*. *Hoxd1* expression in the VPs of trans-heterozygous *HoxD^Del(attP-TpsB2)lac/Del(1-13)^* mutant embryos was completely abolished, in contrast with *Del(1-13)* heterozygous control littermates (Fig. 4B). Conversely, the *HoxD^Del(TpSB2-TpSB3)lac^* allele (hereafter *Del(SB2-SB3)*(Andrey et al., 2013)) carries a large deletion that encompasses the whole T-DOM except the D1 region. We detected robust *Hoxd1* expression in the VPs of E12.5 *HoxD^Del(TpSB2-TpSB3)lac/Del(1-13)^* trans-heterozygous mutant embryos (Fig. 4B).

**Figure 4.**
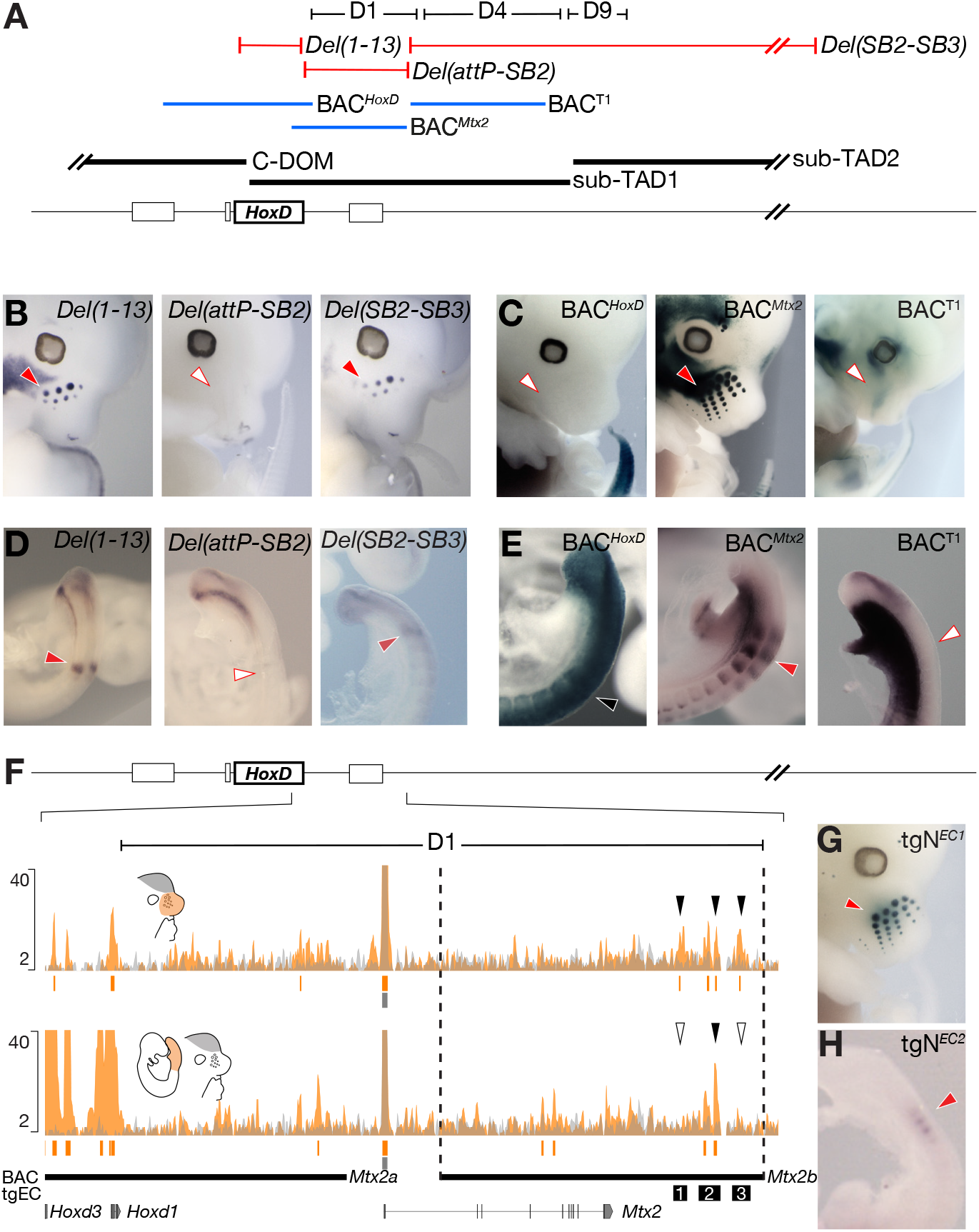
Transcriptional regulation of *Hoxd1.* **A.** The three deletion lines used are shown in red lines and the transgenic BACs in blue lines. The extent of the D1 to D9 regions are shown on top (black) as well as the positions of both sub-TADs below (thick black lines). **B.** WISH using a *Hoxd1* probe on E12.5 mouse embryos. *Hoxd1* mRNAs were detected in both the control and the Del(SB2-SB3) lines (red arrowheads), yet the signal was absent from Del(attP-SB2) mutant embryos (white arrowheads). **C.** X-gal staining on E12.5 mouse embryos carrying randomly integrated BACs. A strong staining was detected in VPs when *BAC^Mtx2^* was used (red arrowhead), whereas staining was not scored with the other two flanking BAC clones (white arrowheads). Therefore, a region directly downstream the *HoxD* cluster is necessary and sufficient to activate *Hoxd1* transcription in VPs, referred to as D1 region as it coincides with this previously defined region. **D.** WISH of *Hoxd1* on E9.5 embryos focusing on tail buds. *Hoxd1* was detected in forming somites in both the control and the *Del(SB2-SB3)* lines (red arrowheads), but was absent from *Del(attP-SB2)* mutant embryos (white arrowhead). **E.** X-gal staining of E9.5 mouse embryos carrying various BAC transgenes. Staining was scored in the entire neural tube and paraxial mesoderm with the BAC^*HoxD*^ (black arrowhead). In contrast, embryos carrying *BAC^Mtx2^* displayed staining in forming somites (red arrowhead), a staining absent when BAC^T1^ was used (white arrowhead). **F.** Enlargement of the D1 region along with a H3K27ac ChIP-seq using dissected E12.5 VPs (orange, top) and E9.5 posterior trunk cells (orange, bottom) superimposed over E12.5 forebrain (FB) cells (grey) (mm10; chr2:74747751-74916570). Above the profiles are the three VP acetylated elements (black arrowheads) and below are MACS2 narrowPeaks for VPs and PTs (orange) and FB (grey). The position of the three EC sequences used as transgenes (tgEC) are shown by numbered black boxes. **G.** X-gal staining in VPs on a E12.5 embryo transgenic for *EC1*. **H.** X-gal staining in forming somites on E9.5 embryo transgenic for the *EC2* sequence.

We complemented these results by producing mouse transgenic embryos carrying bacterial artificial chromosomes (BACs) spanning the *HoxD* cluster and/or parts of sub-TAD1 (in blue in Fig. 4A). In agreement with the analysis of the T-DOM deletions, the BACs covering either the *HoxD* cluster (BAC^*HoxD*^ (Schep et al., 2016) or mapping outside of the D1 region (BAC^*T1*^ (Delpretti et al., 2013) did not display any *lacZ* reporter activity in E12.5 to E13.5 VPs (Fig. 4C). Instead, mouse embryos transgenic for *BAC^Mtx2^* (Allais-Bonnet et al., 2021), which covers most of the D1 region, displayed strong reporter expression in the VPs (Fig. 4C). Noteworthy, expression of *Hoxd1* was also abolished from facial mesenchymal progenitors in *Del(attP-SB2)* mouse embryos, while it was reported by X-gal staining of *BAC^Mtx2^* transgenic embryos, illustrating that the D1 region regulates the transcription of *Hoxd1* in several tissues.

To assess whether the D1 region is also necessary and sufficient for the specific *Hoxd1* expression pattern in somites, we looked at *Hoxd1* transcription in posterior trunk cells using the same set of mutant alleles. *Hoxd1* expression was abolished in developing somites of *Del(attP-SB2)* E9.5 embryos, compared to their control *Del(1-13)* littermates (Fig. 4A, D), while similar transcript levels were maintained in the tailbuds, suggesting that *Hoxd1* regulation in both structures involves different regulatory sequences. Accordingly, embryos transgenic for *BAC^Mtx2^* displayed strong lacZ staining in somites but not in the tailbud, with stronger activity in the last formed somites (Fig. 4A, E) thus closely mimicking native *Hoxd1* transcription in these structures (Zákány et al., 2001). In contrast, *lacZ* expression was uniform along the paraxial mesoderm of BAC^*HoxD*^ transgenic embryos, yet without showing any *Hoxd1*-like specific reporter activity in the forming somites. Altogether, these results confirmed the regulatory potential of the mouse D1 region for *Hoxd1* expression patterns in facial tissues and in forming somites.

### Emergence of tissue-specific enhancers within conserved chromatin domains

*Hoxd1* transcription in VPs could have appeared through the evolution of a new CREs, in association with the emergence of these structures in mammals. An alternative possibility would be the re-deployment of a pre-existing element already at work in somites. The latter hypothesis comes from the fact that in both VPs and somites, transcription of *Hoxd1* follows a cyclic dynamic, with a time progression in phase with the production either of somites or of whisker pads. To address this question, we set up to isolate individual VP enhancers and see if they would be active in somites. We generated two smaller BAC transgenes through homologous recombination of a *lacZ* reporter construct, starting from *BAC^Mtx2^.* While *BAC^Mtx2a^* displayed no signal in VPs, *BAC^Mtx2b^* reported stable β-galactosidase activity (Fig. S4A). Within the *BAC^Mtx2b^* region, three candidate enhancers (ECs) were selected based on H3K27ac enrichment, one of them being also H3K27ac-positive in posterior trunk cells (Fig. 4F, black and white arrowheads). Their regulatory potential was assessed using transient *lacZ* reporter assays in transgenic mouse embryos and we identified a strong and reproducible enhancer activity of *tgN^EC1^* in VPs, comparable to that observed in embryos transgenic for *BAC^Mtx2^* (Fig. 4G). Instead, β-galactosidase activity was not detected neither in *tgN^EC2^,* nor in *tgN^EC3^* transgenic embryos (Fig. S4B). We next tested these candidate enhancers at E9.5 to evaluate their regulatory potential in forming somites. *EC2,* which was the only candidate sequence with significant H3K27ac enrichment in posterior trunk cells, displayed reproducible activity in forming somites (Fig. 4H), unlike *tgN^EC1^* or *tgN^EC3^* (Fig. S4C). We concluded that the only VP enhancer sequence identified was not active during somite formation, unlike a somite enhancer located nearby (2.7 kb), within the D1 region.

The fact that two independent tissue-specific regulations rely on enhancers located in a close proximity within a region of strong interactions suggests that a constrained topology may help the emergence of novel regulatory sequence (Darbellay and Duboule, 2016b). To investigate how CREs might evolve within domains of preferential interactions and how their activity might relate to the observed transcriptional outputs, we compared our mouse and chicken H3K27ac datasets and WISH analyses with the percentage of DNA sequence conservation. The number and distribution of putative CREs (pCREs, defined as non-coding H3K27ac peaks) in the D1, D4 and D9 regions of both mouse and chicken were thus analyzed. Conserved non-coding sequences (CNEs) were annotated by using blastz pairwise alignments downloaded from UCSC and from which all elements overlapping with exons or promoters were discarded (Fig. S4D, E).

Overall, the number and distribution of pCREs in the different regions correlated with the transcriptional activity of *Hoxd* genes. A majority of pCRE sequences did not coincide with CNEs, suggesting a lineage-specific evolution of regulatory elements. Amongst those pCREs coinciding with CNEs, several displayed regulatory activities in one species only (Fig. S4, black arrows) indicating a divergence in the combinations of CREs at work, even in posterior trunk cells where the transcriptional activity is expected to be similar in both species. This observation indicates that the sequence conservation of individual regulatory elements is not an obligatory requirement to maintain similar transcription patterns.

### Divergent enhancer activities of conserved sequences

To take this analysis genome-wide, we compared the H3K27ac profiles across different mouse and chicken embryonic tissues in relation to the presence of CNEs, by adding available proximal and distal forelimb datasets (Beccari et al., 2016; Yakushiji-Kaminatsui et al., 2018) to our posterior trunk and skin H3K27ac ChIP-seq datasets. pCREs were separated into putative enhancers and promoters, hereafter referred to as ‘enhancers’ and ‘promoters’ for simplicity, with ‘enhancers’ mapping more than 2 kb away from a gene transcription start site and ‘promoters’ mapping less than 2 kb away. pCREs which were H3K27ac-positive in more than one of the analyzed tissues were categorized as ‘pleiotropic’ while those being active in only one tissue were named ‘specific’ for the sake of classification. In both mouse and chicken, most promoters were pleiotropic while the majority of enhancers were specific (Fig. 5A), a result in agreement with the usual pleiotropic expression of genes carrying important developmental functions. When enhancers were divided into ‘CNEs’ or ‘non-CNEs’, i.e. displaying or not sequence similarities between mouse and chicken, the proportion of pleiotropic enhancers was higher amongst the former (Fig. 5A; p-value < e-15, Fisher test), an increased level of sequence conservation that may reflect the higher evolutionary pressure on enhancers endorsing multiple regulatory functions.

**Figure 5.**
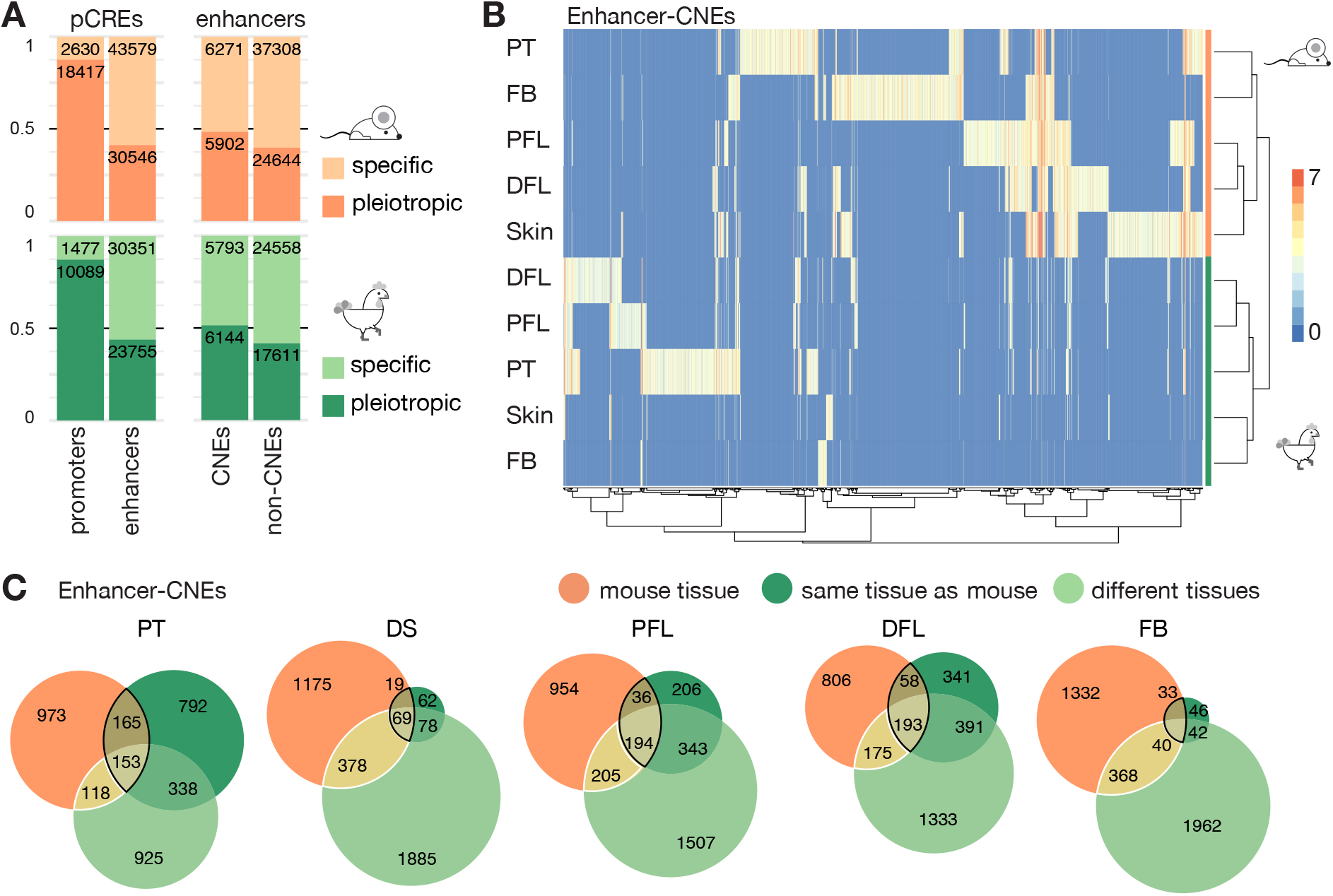
Conserved sequences with divergent functions. **A.** Bar plots showing the proportion of putative *cis*-regulatory elements (pCREs) harbouring the H3K27ac mark in one (‘specific’) *versus* more than one (‘pleiotropic’) tissues, in either mouse (orange) or chicken (green). For simplification purposes, distal pCREs (>2kb away from the transcription start site) are referred as ‘enhancers’ and proximal pCREs (<2kb away from transcription start site) are called ‘promoters’. The different proportions are shown when comparing enhancers and promoters (left), and conserved enhancers to non-conserved enhancers (right). The proportion of specific *versus* pleiotropic elements is shown for CNEs and non-CNEs enhancers (right). In both mouse and chicken, enhancers are more specific than promoters and amongst enhancers, conserved elements are found acetylated in more tissues than non-conserved elements. **B.** Hierarchical clustering obtained using pheatmap R package. On the *X*-axis are shown enhancer-CNEs i.e., conserved sequences, which are non-coding in mouse and in chicken and which overlap with a H3K27ac mark in at least one species. On the *Y*-axis are displayed those tissues used to obtain MACS2 processed peaks. The values on the heatmap correspond to the enrichment scores of the peaks. Enhancer-CNEs cluster by species and not by tissues. **C.** Euler diagrams representing the proportion of enhancer-CNEs, per tissue, acetylated in mouse (orange), in the corresponding chicken tissue (dark green), or in unrelated chicken tissue (light green), showing that conserved sequence diverge in their regulatory activities. The intersection between the same tissue in mouse and in chicken is delineated by a black line, and the intersection between a chicken tissue different from the mouse tissue is delineated by a white line. In **B.** and **C.** PFL, proximal forelimb; DFL, distal forelimb; posterior trunk, posterior trunk; DS, dorsal skin.

Next, we used CNEs to compare the regulatory activities of murine and chicken orthologous sequences. In agreement with what was observed at the *HoxD* locus, the enhancer activities of CNEs, as inferred by H3K27ac peaks, more often diverged than they matched between the related mouse and chicken tissues (Fig. 5B, C). Hierarchical cluster analysis revealed that the enhancer profiles of CNEs were more similar between different tissues of the same species, than between the same tissues in mouse and chicken (Fig. 5B), a surprising observation as these CNEs were initially selected due to their sequence conservation between the two species. We hypothesized that some sequences could have been kept over time despite a divergence in their regulatory activities because they would endorse other functions in other tissues. To evaluate this possibility, we represented on Euler diagrams the relationships between mouse and chicken CNEs displaying enhancer features (Fig. 5C). Each mouse tissue was used as a reference in independent diagrams, where chicken enhancer-CNEs were represented as different sets depending on whether they were active in the same or in different tissues when compared to the mouse reference.

The intersection between the same tissue in mouse and in chicken represents elements that have both a conserved sequence and regulatory function (Fig. 5C, overlaps circled in black). This intersection is smaller than any difference between two sets, which indicates that DNA sequence similarity between two species can be maintained amongst regulatory sequences despite divergences in their regulatory functions. We could observe that CNEs often display enhancer activities in a chicken tissue that is different from the mouse tissue, supporting this conclusion (Fig. 5C, overlaps circled in white). As a consequence, the regulatory activity of a given sequence conserved between these two species cannot be used to predict its function(s) in the two species.

Altogether, we show that both the global genomic organisation of the *<u>HoxD</u>* locus and specific domains of preferential enhancer-promoter interactions are highly conserved between mouse and chicken. However, the regulatory activities of individual CREs are largely species-specific and such sequences are systematically found within those segments of regulatory landscapes that preferentially interact with the promoters of the most highly transcribed genes. We also show that the lineage-specific evolution of regulatory activities could either generate new regulatory relationships or, conversely, maintain ancestral transcription patterns.

## DISCUSSION

### Versatile *Hoxd* gene functions in skin appendages

*Hox* genes play a dual role in skin appendage development. While their colinear expression is initially necessary to pattern the skin dermis (Kanzler et al., 1997), they are subsequently transcribed in the dermal papillae and/or ectoderm of embryonic and adult hair pelage follicles and vibrissae precursors (Godwin and Capecchi, 1998; Packer et al., 2000; Reynolds et al., 1995; Yu et al., 2018), as well as in the avian buds (Chuong et al., 1990; Kanzler et al., 1997). Here we show that the dermal papillae of embryonic mouse vibrissae express *Hoxd1* to *Hoxd4,* with the former gene’s mRNAs being the first detected, followed by considerably lower levels of *Hoxd3* and *Hoxd4* RNAs. The transcription in VPs is driven by CREs located in a region encompassing approximately 120kb of DNA adjacent to the telomeric end of the mouse *HoxD* cluster, a region that is preferentially contacted by the *Hoxd1* gene. Therefore, a proximity effect is observed in the building of a general chromatin architecture, based on constitutive contacts between the *Hoxd1* region and the neighboring sub-TAD, whereby most of the *Hoxd1* regulatory sequences are located in the adjacent portion of the regulatory landscape.

While a detailed functional characterization of *Hoxd* genes during the development of whiskers was not the aim of this work, we did not observe any major morphological alteration in the vibrissae of mice carrying genetic deletions of this precise region, despite the complete abrogation in the expression of *Hoxd1* to *Hoxd4* in these structures. This lack of any visible abnormal phenotype may be due to functional redundancy with other *Hox* paralogy groups expressed there, such as *Hoxc8* (Yu et al., 2018), to a lack of analytical power or of the appropriate behavioral paradigms that may have revealed functionally abnormal whiskers. The diverse expression of *Hox* paralogs in mammalian and avian skin appendages nonetheless illustrates the plasticity in the usage of *Hox* gene functions in these structures. This plasticity likely relies upon the independent acquisition of specific CREs for each gene cluster, as these skin derivatives emerged after the genome duplication events thought to be at the origin of the vertebrate lineage (Holland et al., 2008; Holland and Garcia-Fernàndez, 1996).

While *Hoxd9, Hoxd11* and *Hoxd13* are expressed in the ectoderm-derived hair fibers (Godwin and Capecchi, 1998; Packer et al., 2000; Reynolds et al., 1995; Yu et al., 2018), we did not observe any *Hoxd1* in the vibrissae shaft, nor any lacZ activity when the *TgBAC^Mtx2^* (carrying the D1 region) transgenics were used, indicating that these expression patterns may depend on different sets of CREs. Our data also show that the expression of *Hoxd3* to *Hoxd9* in the ectoderm of the chicken FPs depends on CREs unrelated to those operating in the mouse VPs. These observations may indicate that different sets of *Hoxd* genes were independently co-opted during the evolution of skin derivatives, likely through *de novo* enhancer acquisition in the mammalian and avian lineages, in a way related to the regulation of *Hoxd9 and Hoxc13* in the embryonic mammary buds and in the nails and hairs of mammals, respectively (Fernandez-Guerrero et al., 2020; Schep et al., 2016). Alternatively, the rapid pace of tegument evolution in tetrapods may have been accompanied by a divergence in enhancer sequences, which makes comparisons difficult. In this context, we can not rule out the possibility that *Hoxd* gene regulation in skin appendages of both mammals and birds derives from an initial pan-*Hoxd* expression pattern both in the ectoderm and mesoderm of skin appendages, which then evolved differently in each lineage through modification of the CRE complement located within the adjacent TAD. The fact that two CNEs acetylated in mouse VPs and chick FPs are also acetylated in mouse and chick posterior trunk cells may indicate that the former structures co-opted parts of the regulatory mechanisms at work in the latter. In this view, DNA sequence conservation may be due to constraints imposed by the function in the trunk rather than by the function in the skin.

### Enhancer acquisition in evolving tetrapod skin appendages

TADs containing pleiotropic genes of key importance for vertebrate development are frequently conserved across tetrapods and topological changes in their organization often correlate with important modifications in gene expression (Eres et al., 2019; Liao et al., 2021; Torosin et al., 2020; Yakushiji-Kaminatsui et al., 2018). Our results show that despite a considerable divergence in the non-coding elements localized in the mouse and chicken T-DOM, the global pattern of interactions is conserved leading to similar chromatin topologies. Indeed, in both cases, different *Hoxd* genes interact preferentially with distinct portions of sub-TAD1. These contacts are for the most constitutive, even though we observed an activity-dependent increase in interactions in the mouse VPs, and in mouse and chicken posterior trunk cells. Of interest, the regions within T-DOM, which were preferentially contacted by the *Hoxd1, Hoxd4* or *Hoxd9* genes (D1 to D9) are organized in a linear sequence opposite to the order of the genes themselves, as if *Hoxd1* was folding on its closest possible chromatin environment within T-DOM, bringing along *Hoxd3* towards the next chromatin segment and positioning the interactions involving *Hoxd9* further away into T-DOM. This sequential positioning of interactions in mirror with spatial colinearity may impose a generic contact pattern, whereby *Hoxd9*, for example, is unable to establish strong interactions with the D1 region and its enhancers, for this region is primarily contacting *Hoxd1,* thus imposing constraints upon the distributions of interactions (see below).

Furthermore, the distribution of H3K27ac-positive regions in the chicken FPs and mouse VPs across these interacting regions correlate well with the specific subsets of *Hoxd* genes expressed in each of these tissues. Finally, the regulation of *Hoxd1* in developing somites relies on evolutionary conserved elements located in the region D1. This general regulatory topology is close to that observed in teleost fishes (Acemel et al., 2016; Woltering et al., 2014), suggesting that it was already in place at the root of the vertebrate lineage and that its hijacking by nascent enhancer sequences, may have favoured the co-option of specific *Hoxd* gene subsets as targets of the novel regulations. The regulation of *Hoxd* genes in the mouse VPs and chicken FPs, as well in the mouse mammary buds (Schep et al., 2016) may illustrate this process.

### Plasticity of the *cis*-regulatory code *versus* conservation in TAD organization

Due to their functional relevance in the regulation of gene expression, CREs tend to be evolutionarily conserved across species (Sandelin et al., 2004; Sanges et al., 2013, 2006). However, several studies have shown that enhancers controlling developmental gene expression can be functionally conserved even across distant phylogenetic lineages despite a strong divergence in their DNA sequences (Ambrosino et al., 2019; Berthelot et al., 2018; Wong et al., 2020) (Eichenlaub and Ettwiller, 2011; Schmidt et al., 2010; Villar et al., 2015). Our inter-species comparison of CNEs enriched in H3K27ac across different embryonic structures revealed that a considerable portion of evolutionary conserved CREs diverged in their patterns of activity between the two species analyzed, in agreement with previous observations (Dermitzakis and Clark, 2002; Schmidt et al., 2010; Vierstra et al., 2014; Villar et al., 2015). This dichotomy between the conservation of enhancer sequences and their function is illustrated by the CREs controlling *Hoxd1* expression in the mouse and chicken developing somites. While some of the sequences active in the mouse posterior trunk are evolutionary and functionally conserved in chicken, such as EC2, which can drive lacZ expression in the last formed somites of the murine embryo, other elements conserved at the nucleotide level displayed H3K27ac enrichment only in the mouse.

In fact, a large fraction of H3K27ac-positive sequences detected either in the mouse or in the chicken posterior trunk cells were not conserved between the two species, suggesting that they evolved in a species-specific manner. In contrast, the high proportion of chicken promoters that display similar H3K27ac enrichments in mice, together with the similar expression patterns observed for mouse and chicken *Hoxd* genes in embryonic trunk cells, demonstrate that conserved gene expression can occur despite a high evolutionary plasticity in the *cis*-regulatory code usage, in agreement with previous reports (Berthelot et al., 2018; Snetkova et al., 2021). This may point to a regulatory strategy where multiple enhancers act coordinately to regulate the same target gene(s) thus creating a quantitative effect in a way similar to the reported action of super-enhancers (Sabari et al., 2018), yet where various subsets of CREs may display distinct tissue-specificities. This process may be implemented preferentially within large regulatory landscapes, as e.g. in the case of *Fgf8* (Marinic et al., 2013) or the TAD controlling *Hoxd* gene transcription in external genitals (Amândio et al., 2020).

### A regulatory playground with constraints

In those cases that were investigated, the respective positions of regulatory sequences within large syntenic regulatory landscapes are generally conserved, and the global regulatory architecture found at one particular developmental locus in one species can be extrapolated to its cognate locus in another amniote species (e.g. (Kragesteen et al., 2018), implying that enhancer-promoter interactions are also evolutionary well conserved, even for CREs acting over long distances (Irimia et al., 2012; Laverré et al., 2021), although some mechanistic elements may vary (Ushiki et al., 2021). In agreement with this, and contrasting with the apparent divergence in the *cis*-regulatory code controlling *Hoxd* gene expression in mouse and chicken embryos, our phylogenetic footprinting and epigenetic analysis revealed that non-coding elements conserved in their DNA sequence between the mouse and chicken genomes were distributed across the two species in a very similar manner. Therefore, the potential gain and/or loss in the two lineages of particular CREs did not significantly impact the internal organization of those conserved regulatory elements, nor did they impact upon the global chromatin topology of the locus. This confirms the observation that even the significant divergence in CREs located at the *HoxD* locus, which accompanied the evolution of the snake body plan, did not result in any substantial alteration of the overall corn-snake *Hoxd* gene interaction map (Guerreiro et al., 2016). Altogether, these observations suggest that the internal organization of the TADs at the *HoxD* locus and the specific interaction profiles for various subsets of *Hoxd* target genes are resilient to considerable variation in their CREs.

These results also highlight the somewhat dual properties that large regulatory landscapes may have imposed in the course of regulatory evolution. On the one hand, a pre-established chromatin topology, as materialized by TADs, may have provided unique ‘structural niches’ to evolve new enhancer sequences, due to the proximity of factors already at work. This might especially apply to landscapes where a strong quantitative parameter may be instrumental, and *a fortiori* when the target gene(s) are acting in combination, coding for proteins of rather low specificities (Bolt and Duboule, 2020). This may be illustrated nowadays by large landscapes, where all kinds of enhancer sequences are mixed within the same TAD. On the other hand, this regulatory playground, while stimulating the emergence of regulations, will constrain their realms of action due both to the very architecture that favors their evolution and to the accessibility to ‘a’ subset of contiguous target genes as a result of the global distribution of interactions at this precise place of the landscape.

## MATERIAL AND METHODS

### Mouse strains and chicken eggs

All mutant strains are listed in Table S1 and were backcrossed continuously to Bl6/CBA mixed animals and maintained as heterozygous stocks. Chick embryos from a White Leghorn strain were incubated at 37.5°C and staged according to Hamburger and Hamilton (1951).

### Enhancer cloning and transgenic animals

The different CRE candidate sequences were amplified using PCR-specific primers (see Table S2) and the Expand High Fidelity PCR system (Roche). Amplified fragments were gel-purified and cloned into the pSK-lacZ vector as in (Beccari et al., 2016). For the transgenesis assays, each lacZ reporter construct was digested with NotI and KpnI and the restriction fragment encoding the potential enhancer sequence, the beta-globin minimal promoter and the lacZ reporter gene was gel-purified and injected into the male pronucleus of fertilized oocytes. F0 embryos were dissected either at E10, or at E12, fixed and stained for lacZ activity according to standard protocols.

### *In situ* hybridization

The probes used in this study for *in situ* hybridization were either previously reported, or produced by PCR amplification of a fragment subsequently ligated in pGEMT Easy vector (Promega). Primers used to amplify the fragment of interest are listed in Table S2. Digoxigenin (Dig) labeled probes were synthesized *in vitro* by linearizing the respective plasmids using specific restriction enzymes and transcribed *in vitro* either with T7/T3, or Sp6 RNA polymerase (Table S3) and the Dig-RNA labeling mix (Roche). The probes were purified using the RNA easy mini kit. WISH experiments were performed as described in (Woltering et al., 2009). Probes are listed in (Table S3). *In situ* experiments in sections were performed according to (Sockanathan and Sockanathan, 2015). Cryostat sections were prepared after fixing embryos in 4% PFA solution at 4°C overnight. Subsequently, embryos were washed three times with PBS and passed to increasing sucrose solutions at 4°C in agitation (15% sucrose solution for 3-4 hours, followed by incubation in 30% sucrose solution overnight) and finally included in OCT blocks. Sections of 20 μm were obtained with a Leica cryostat and stored at −80°C.

### ChIP-seq

For genomics analyses, we used the mouse assembly mm10 and the chicken galGal6 and we plotted genomic data with pyGenomeTracks 3.6 (Lopez-Delisle et al., 2021). ChIP-seq experiments were performed using the ChIP-IT High sensitivity kit (active motif according to manufacturer instructions with minor modifications). For the mouse samples, 12 pairs of E12.5 WPs, or one hundred E8.75 posterior trunks were used, the latter dissected at the level of the 2^nd^ to 4^th^ pair of somites. For the chicken samples, the dorsal skin of five HH35 embryos or the posterior trunk or 80 HH19-21 posterior trunk regions were isolated. In all cases, the samples were fixed for 10 min in 1% formaldehyde at RT and the crosslinking reaction was quenched with Glycine. Subsequently, nuclei were extracted and sonicated in (Tris HCl pH 8.0 50mM, EDTA 10mM, 1% SDS) to an average fragment size of 200-500bp using a Diagenonode Bioruptor Sonicator. Ca. 10 to 20 μg of sonicated chromatin were diluted tenfold with Dilution buffer (HEPES pH 7.3 20mM, EDTA 1mM, NaCl 150mM, NP40 0.1%) and incubated with 2 μg of anti-H3K27ac antibody (Abcam, ab4729). All buffers contained 1X Complete Proteinase Inhibitor cocktail (EDTA-free; Roche) and 10mM Sodium Butyrate to prevent protein degradation and Histone de-acetylation. Chromatin–antibody complexes were immunoprecipitated with Protein G-coupled agarose beads-purified and de-crosslinked according to manufacturer instructions. Approx 5 to 10 ng of purified DNA were used for ChIP library preparation by the Geneva IGE3 Genomics Platform (University of Geneva) and sequenced to obtain 100-bp single-end reads on an Illumina HiSeq2500 or HiSeq4000 system. Published fastq files were downloaded from SRA for H3K27ac: mouse DFL (GSM2713703, <u>SRR5855214</u>), mouse PFL (GSM2713704, <u>SRR5855215</u>), chicken DFL (GSM3182462, <u>SRR7288104</u>), chicken PFL (GSM3182459, <u>SRR7288101</u>); and for CTCF in mouse PT (GSM5501395, <u>SRR15338248</u>). ChIP-seq reads processing was done as in (Amândio et al., 2020) with some modifications, using the laboratory Galaxy server (Afgan et al., 2016). Adapters and bad-quality bases were removed with Cutadapt (v1.16.8) (Martin, 2011) (options -m 15 -q 30 -a GATCGGAAGAGCACACGTCTGAACTCCAGTCAC). Reads were mapped to the mouse genome (mm10) and to the chicken genome (galGla6) using Bowtie2 (v2.4.2) (Langmead and Salzberg, 2012) with standard settings. Only alignments with a mapping quality >30 were kept (Samtools v1.13) (Danecek et al. 2021). H3K27ac ChIP mapped reads were down sampled with Samtools view (v1.13) according to the sample with the smallest number of reads after filtering and duplicate removal (mouse: 71741225 reads; chicken 25872853 reads). Mouse CTCF ChIP mapped reads were down sampled with a fraction of 1/4 and chicken CTCF ChIP reads were not down sampled. When two replicates were available, alignment bam files were merged with Samtools merge (v1.13) and subsampled to the same number as single replicates. The coverage and peaks were obtained as the output of MACS2 (v2.1.1.20160309.6) (Zhang et al., 2008) with a fixed extension of 200 bp (--call-summits --nomodel --extsize 200 -B), and, for the WP sample, the q-value cutoff for peak detection was set to 0.1 instead of the 0.05 default. CTCF motif orientation was assessed using peaks extended by 100 bp each side and the CTCFBSDB 2.0 database (Ziebarth et al., 2012) with MIT_LM7 identified motifs.

### MAR annotation

MARs are genomic segments of the T-DOM with the highest density of specific H3K27ac coverage and are annotated as follow: For mm10 chr2:74872605-75696338 and for galGal6 chr7:15856252-16252401, sliding windows are of 10 kb every 2 kb. For each window the coverage of the ChIP was computed and the difference with the coverage in the corresponding H3K27ac ChIP in the brain was considered as the specific signal. The smallest sequence of consecutive windows cumulating 30% (for the DS and WP samples) or 50% (for the PT samples) of the total specific signal is the MAR. See ChIP_scanRegionForHighestDensity_2files.py at https://gitlab.unige.ch/Aurelie.Hintermann/hintermannetal2022 for all details.

### pCREs

pCREs are defined as genomic intervals that overlap with H3K27ac Macs2 peaks in at least one of the samples analyzed per species. To obtain the genomic coordinates of pCREs, Macs2 peaks of all samples were pooled into one file, the size of each peak was increased by 660 bp each side to account for nucleosome occupancy with bedtools slop (v2.30.0), and overlapping Macs2 peaks were combined into a single pCRE interval using bedtools merge. pCREs were annotated according to their colocalization with tissue-specific H3K27ac peaks, exons and sequence conservation using bedtools intersect. pCREs were quantified using R (v3.6.1) and the bar plots of Figure 5A were done with the ggplot2 package (v3.2.1). See CNEs_01.sh at https://gitlab.unige.ch/Aurelie.Hintermann/hintermannetal2022 for all details.

### CNEs

Pairwise blastz alignments were downloaded from http://hgdownload.soe.ucsc.edu/goldenPath/mm10/vsGalGal6/mm10.galGal6.synNet.maf.gz and transformed to intervals with maf to interval (Blankenberg et al., 2011). Conserved elements were considered non-coding if they would not overlap with annotated exons neither in mouse nor in chicken. The synteny plot on Figure S3A was done using the R package ggplot2 (v3.2.1). CNEs were annotated according to their colocalization with tissue-specific H3K27ac peaks, exons and sequence conservation using bedtools intersect. pCREs were quantified using R (v3.6.1) and the bar plots of Figure 5A were done with the ggplot2 package (v3.2.1) (R Core Team (2019). R: A language and environment for statistical computing. R Foundation for Statistical Computing, Vienna, Austria. URL https://www.R-project.org/.). The heatmap in Figure 5B was done using the pheatmap R package (v1.0.12) using a matrix which columns are CNEs that overlapped with an enhancer in at least one tissue in mouse or in chicken, rows are the tissues in which acetylation peaks were called and values are the scores of acetylation peaks. If a CNE comprised several peaks of the same tissue, the average was done. In Figure 5C, the relationships of the three sets formed by the conditions of being acetylated in the mouse tissue, in the chicken tissue or in a chicken tissue excepted the one considered were represented on Euler diagramms using the eulerr R package (v6.0.0), considering independently each tissue analyzed (i.e. PT DFL PFL Skin and FB). See CNEs_01.sh at https://gitlab.unige.ch/Aurelie.Hintermann/hintermannetal2022 for all details.

### 4C-seq experiments

4C-seq experiments were performed according to (Noordermeer et al., 2011). Briefly, the tissues were dissected, dissociated with collagenase (Sigma Aldrich/Fluka) and filtered through a 35 microns mesh. Cells were fixed with 2% formaldehyde (in PBS/10%FBS) for 10 min at RT and the reaction was quenched on ice with Glycine. Nuclei were extracted using a cell lysis buffer (Tris pH 7.5 10 mM, NaCl 10 mM, MgCl2 5 mM, EDTA 0.1 mM,) and stored at -80°C. Approximately 200 E9.5 posterior trunk regions dissected at the level of the 2^nd^ to 4^th^ pair of somites, as well as 120 chicken HH19-21 posterior trunk samples, were used. Nuclei were digested with NlaIII (New England Biolabs) and ligated with T4 DNA ligase HC (Promega) in diluted conditions to promote intramolecular ligation. Samples were digested again with DpnII (New England Biolabs) and re-ligated with T4 DNA ligase HC. These templates were amplified using Expand long template (Roche) and PCR barcoded primers flanked with adaptors. For each library, 8 to 10 independent PCR reactions were pooled together and purified using the PCR purification kit (Qiagen). Multiplexed libraries were sequenced on Illumina HiSeq 2500 to obtain 100 bp single-end reads. Demultiplexing, mapping and 4C-seq analysis were performed using a local version of the pipeline described in (David et al., 2014), on the mouse mm10 and chicken galGal6 genome assemblies. The profiles were smoothened using a window size of 11 fragments and normalized to the mean score in +/-5 Mb around the viewpoint. When multiple independent biological replicates were available, average 4C-seq profiles were calculated. When viewpoints were subtracted, the score of each window of the normalized, smoothed profiles were subtracted (See 4C_subset_subtract at https://gitlab.unige.ch/Aurelie.Hintermann/hintermannetal2022) for all details.

### Capture Hi-C-seq experiments

The SureSelectXT RNA probe design and capture Hi-C experiments were performed as described in (Bolt et al., 2021b) for mouse and as in (Yakushiji-Kaminatsui et al., 2018) for chicken. Dissected tissues were processed as in (Yakushiji-Kaminatsui et al., 2018) with the following changes: cells were cross-linked in 2% formaldehyde/PBS and washed three times instead of being quenched by glycine. Hi-C-libraries were prepared as in (Yakushiji-Kaminatsui et al., 2018). The first part of the data analysis was performed on our local galaxy server (Afgan et al. 2016). Raw reads were preprocessed with CutAdapt (v1.16) (-a AGATCGGAAGAGCACACGTCTGAACTCCAGTCAC -A AGATCGGAAGAGCGTCGTGTAGGGAAAGAGTGTAGATCTCGGTGGTCGCCGTATCATT -- minimum-length=15 --pair-filter=any --quality-cutoff=30) (Martin 2011). Then, Hicup (v0.6.1) (Langmead and Salzberg 2012; Dryden et al. 2014) and Samtools 1.2 (Danecek et al. 2021) were used with default parameters. The pairs were then loaded to 5-kb (mouse) and 2.5-kb (chicken) resolution matrices with Cooler v0.7.4 (Abdennur and Mirny 2020). These matrices were balanced with HiCExplorer hicCorrectMatrix (v3.7.2) using ICE as correction method, the mad threshold from HiCExplorer hicCorrectMatrix diagnostic plot as minimum threshold value and 5 as maximum threshold value. The TAD separation scores were computed with HiCExplorer hicFindTADs. (Ramírez et al. 2018; Wolff et al. 2020) with a fixed window of 240 kb in the mouse and 120 kb in the chicken. See CHiC_processRaws.sh https://gitlab.unige.ch/Aurelie.Hintermann/hintermannetal2022 for all details.

## ETHICS APPROVAL

All experiments involving animals were performed in agreement with the Swiss Law on Animal Protection (LPA), under license No GE 81/14 (to DD).

## DATA AVAILABILITY

All raw and processed datasets are available in the Gene Expression Omnibus (GEO) repository under accession number GSE195592. All scripts necessary to reproduce figures from raw data are available at https://gitlab.unige.ch/Aurelie.Hintermann/hintermannetal2022.

## AUTHORS CONTRIBUTIONS

A. H. : Designed experiments, realized experiments and interpreted the results, wrote the manuscript

I. G. : Did some cloning and *in situ* hybridizations

C. C. B. : Provided unpublished datasets and helped for some the CHi-C experiments

L. L.-D.: Helped with the bioinformatic analyses

S. G. : Genotyped and helped analyze mouse mutants.

D. D. : Designed experiments, interpreted some results and wrote the paper.

L. B. : Designed experiments, interpreted some results and wrote the paper.

## ACKNOWLEDGEMENTS

We thank all members of the Duboule laboratories for comments and discussions and Gregory Loichot for artwork. This work was supported in part by using the resources and services of both the Gene Expression Core Facility at the School of Life Sciences of the EPFL in Lausanne and the genomic platform from the IGE3 centre of the University of Geneva. Transgene injections were performed at the transgenesis platform of the University of Geneva Medical School.

## LEGENDS TO SUPPLEMENTARY FIGURES

**Supplementary Figure S1.**
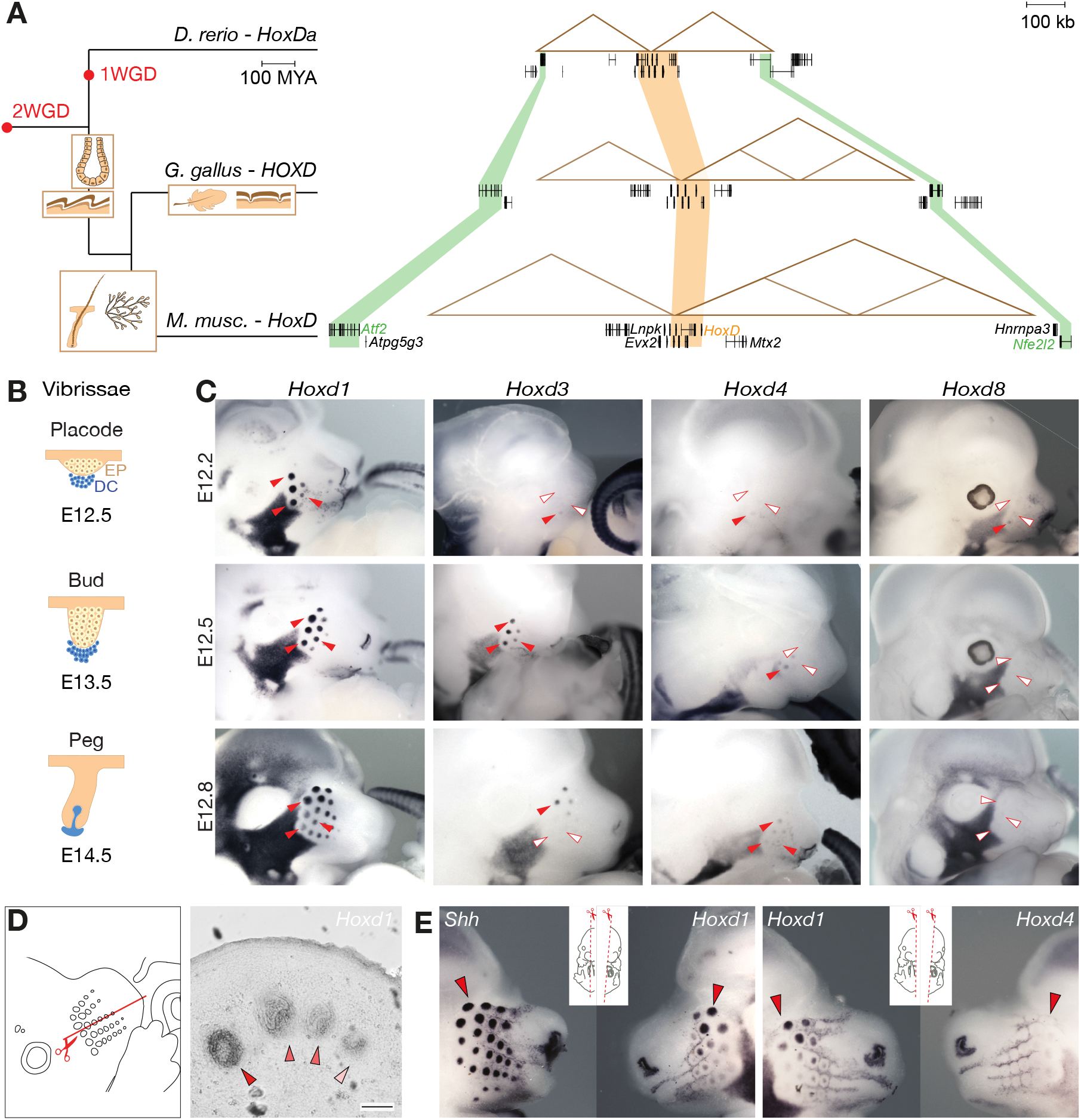
Evolution of the *HoxD* locus and expression of *Hoxd* genes in the developing VPs. **A.** On the left is a schematic illustration of a vertebrate phylogenetic tree showing the emergence of some skin derivatives (glands, scales, feathers, hairs). On the right are the corresponding versions of the *HoxD* locus, with the orthologous *HoxD* gene clusters as shaded orange areas and other orthologous genes shown in green to help delimit the extent of the conserved regulatory landscapes. **B.** Schematic representation of vibrissae morphogenesis: epithelial placode (EP, yellow cells); dermal condensation (DC, blue cells). The approximate developmental stage is indicated below, showing the delay between vibrissae and hair morphogenesis. The dorsal skin of the upper back was used as reference for hairs. **C.** WISH on three successive E12 staged embryos (indicated on the left) using four *Hoxd* genes as probes (top). The same three placodes are pointed by red (positive) or white (negative) arrowheads. **D.** Schematic of the transversal section on E13.5 VPs (left), analyzed in ISH with a *Hoxd1* probe (right). The scale bar in the right picture corresponds to 100 μm and the *Hoxd1* stained VPs at peg stage (see below) are indicated by red arrowheads of decreasing intensities from left to right. The staining is localized in dermal cells. **E.** WISH on E12.5 bisected mouse embryos. *Hoxd1* mRNAs were detected after those for *Shh* (left) and before *Hoxd4* mRNAs (right).

**Supplementary Figure S2.**
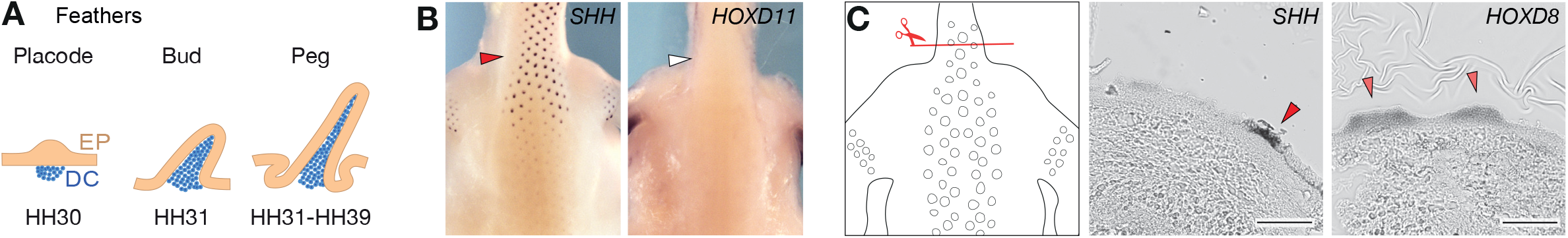
Feather development and transcription of *Hoxd* genes. **A.** Schematic of feather development between stage HH20 and HH35 with ectodermal placode (EP, yellow) and dermal condensation cells (DC, blue). **B.** WISH on HH35 chicken embryos, dorsal view of the upper back, with arrowheads pointing toward the neck at the shoulder level. *HOXD11* transcripts were not detected in FPs (right), unlike those of SHH (left). **C**. Transversal section through a HH35 chicken skin of the upper back (red), corresponding to the two sections stained by ISH on the right. The scale bar is 100 μm. *SHH* is used as marker of the epithelial placode (red arrowhead). *HOXD8* is also transcribed in epithelial placodes, matching -or slightly larger than- the *SHH* domain.

**Supplementary Figure S3.**
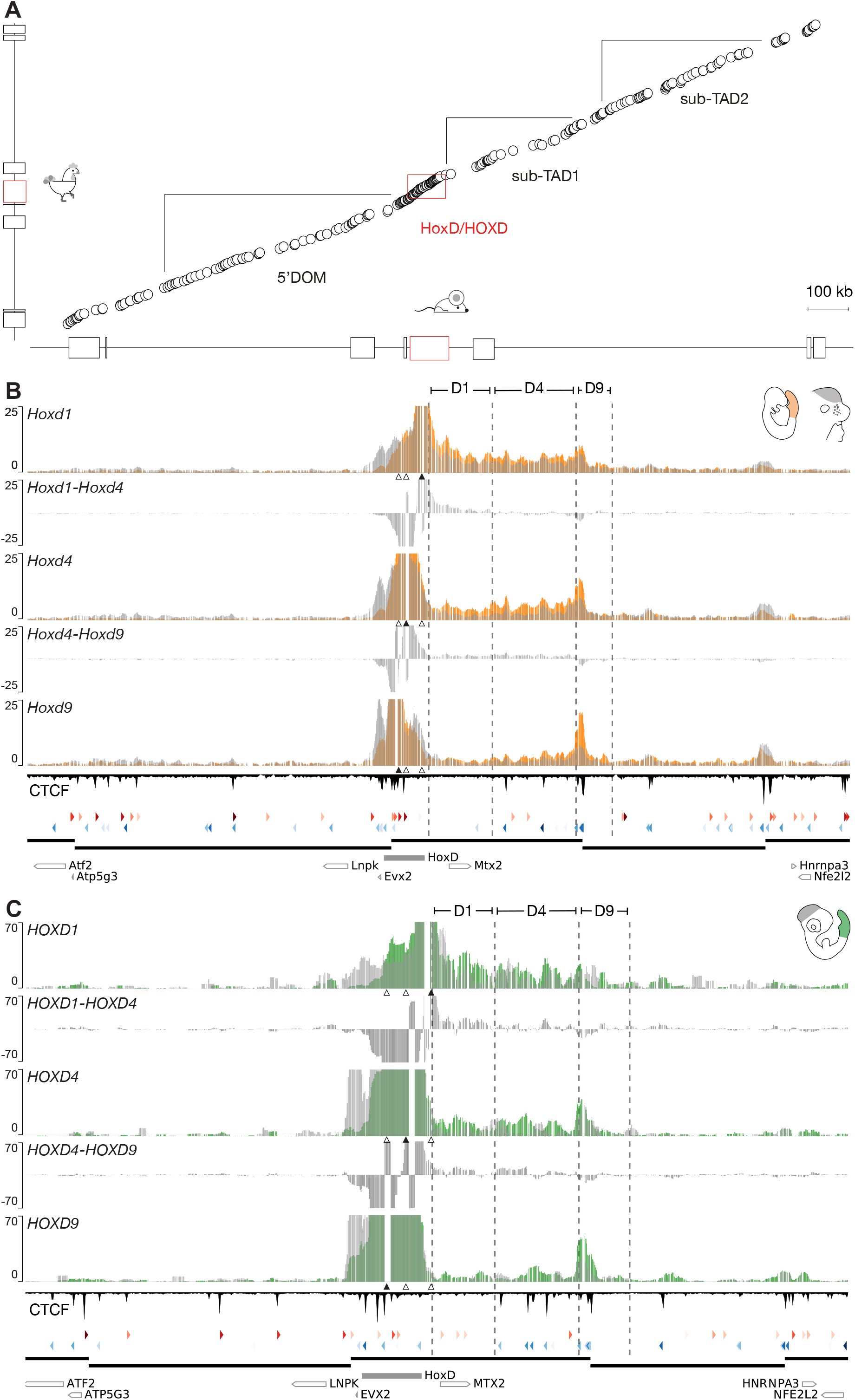
Comparative 4C-seq analyses at the mouse and chicken *HoxD* loci. **A.** Synteny plots representing sequences conserved between the mouse and the chicken *HoxD* loci. On the X-axis is the mouse locus (mm10, chr2:73800000-75800000) and on the Y axis is the chicken locus (galGal6, chr7:15790000-16700000, inverted x-axis). The scale is the same for both axes. TADs and sub-TADs are indicated by black lines and the *HoxD* cluster by a red rectangle. Schematic representations of the locus are on top of each axis. Despite a mouse locus that is in average 2.2 times larger than its chicken counterpart, the order of the conserved sequences is maintained, showing the absence of substantial genomic rearrangement at this locus. **B**. On the Y axis are 4C-seq normalised scores per feature, with E9.5 mouse posterior trunk cells scores (orange) superimposed over E12.5 FB cells (grey). Below the 4C tracks, the positions of the *Hoxd1, Hoxd4* and *Hoxd9* viewpoints are represented by triangles, with a black triangle for the viewpoint used in the track. Between each track, the subtraction of the two adjacent viewpoints is represented (black). The preferentially interacting regions D1, D4 and D9 were annotated manually and, to facilitate comparisons between their positions in the mouse and the chicken locus, orthologous CNEs located close to the edge of each region D are shown (vertical dashed lines). CTCF ChIP-seq profiles are displayed below the 4C tracks, with red and blue arrowheads below indicating their orientation. Color intensity is proportional to motif score. Data mapped on mm10, chr2:73800000-75800000. **C.** On the Y axis are 4C-seq normalised scores per feature, using HH18 chicken posterior trunk cells (green), superimposed with the profile obtained with HH18 brain cells (grey). Data shown for *HOXD1* (top) *HOXD4* (middle) and *HOXD9* (bottom) viewpoints, with a black arrowhead indicating the position of the viewpoint. Reads mapped on galGal6, chr7:15790000-16700000, inverted x-axis.

**Supplementary Figure S4.**
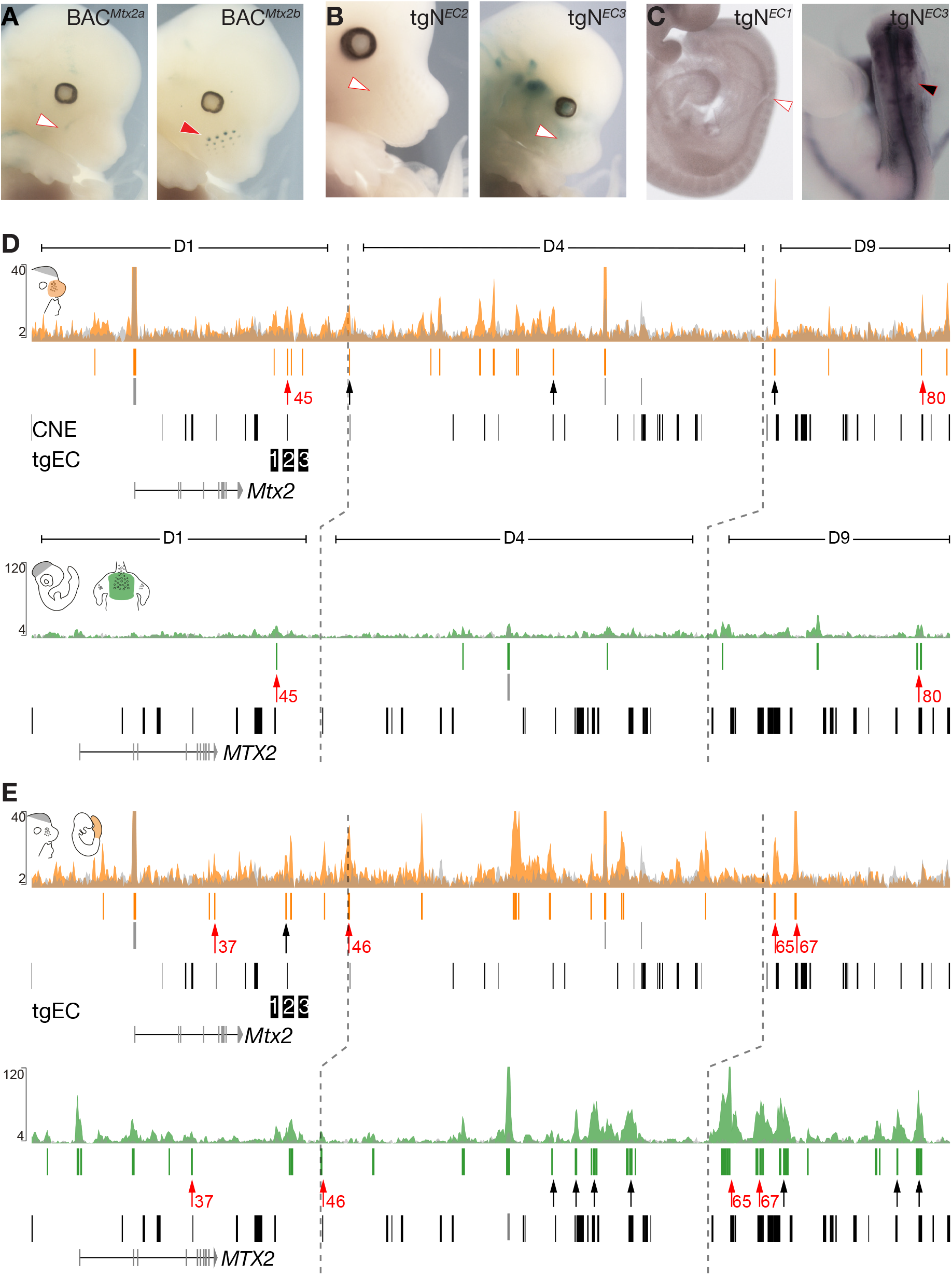
Comparison of conserved and non-conserved CREs in mouse and chicken embryonic tissues. **A.** X-gal staining of E12.5 transgenic embryos carrying either the first half (left) or the second half (right) of *BAC^Mtx2^.* The entire *BAC^Mtx2^* recapitulated the pattern of expression of *Hoxd1* in VPs. **B.** Staining of embryos carrying either the *Tg^EC2^* or the *TgN^EC3^* transgenes, which show no regulatory activity in VPs at E12.5. **C.** Staining of E9.5 mouse embryos carrying the *TgN^EC1^* or the *TgN^EC3^* reporter constructs, which show the absence of signal in forming somites. **D.** H3K27ac ChIP-seq profiles magnifications over the mouse and chicken sub-TAD1. Top: Mouse E12.5 dissected VPs (orange) along with the mouse E12.5 forebrain cells track (grey)(mm10, chr2:74775737-75222876). Below is the profile for the chicken HH35 dorsal skin (green), superimposed to HH18 brain cells (grey) (galGal6, chr7:16033612-16252401, inverted x-axis). Processed MACS2 narrowPeaks are in orange/green vertical lines and blastz conserved sequences in black vertical lines, below each track. The peaks overlapping CNEs are pointed by black arrows. The peaks corresponding to CNEs in both species and in equivalent tissues are pointed by red arrows. Vertical dashed lines represent the position of corresponding CNEs at the edge of each region **E.** H3K27ac ChIP-seq focusing on sub-TAD1. Top: The profile for mouse E9.5 posterior trunk cells (orange) are superimposed to mouse E12.5 forebrain cells (grey). Bottom: Profiles of chicken HH18 embryonic posterior trunk cells (green) and of HH18 brain cells.

## LEGENDS TO SUPPLEMENTARY TABLES

**Supplementary Table S1.** List of mouse lines

**Supplementary Table S2.** Primers used to clone candidate enhancer sequences

**Supplementary Table S3.** Probes for *in situ* hybridization

---

**Supplementary Table 1.**
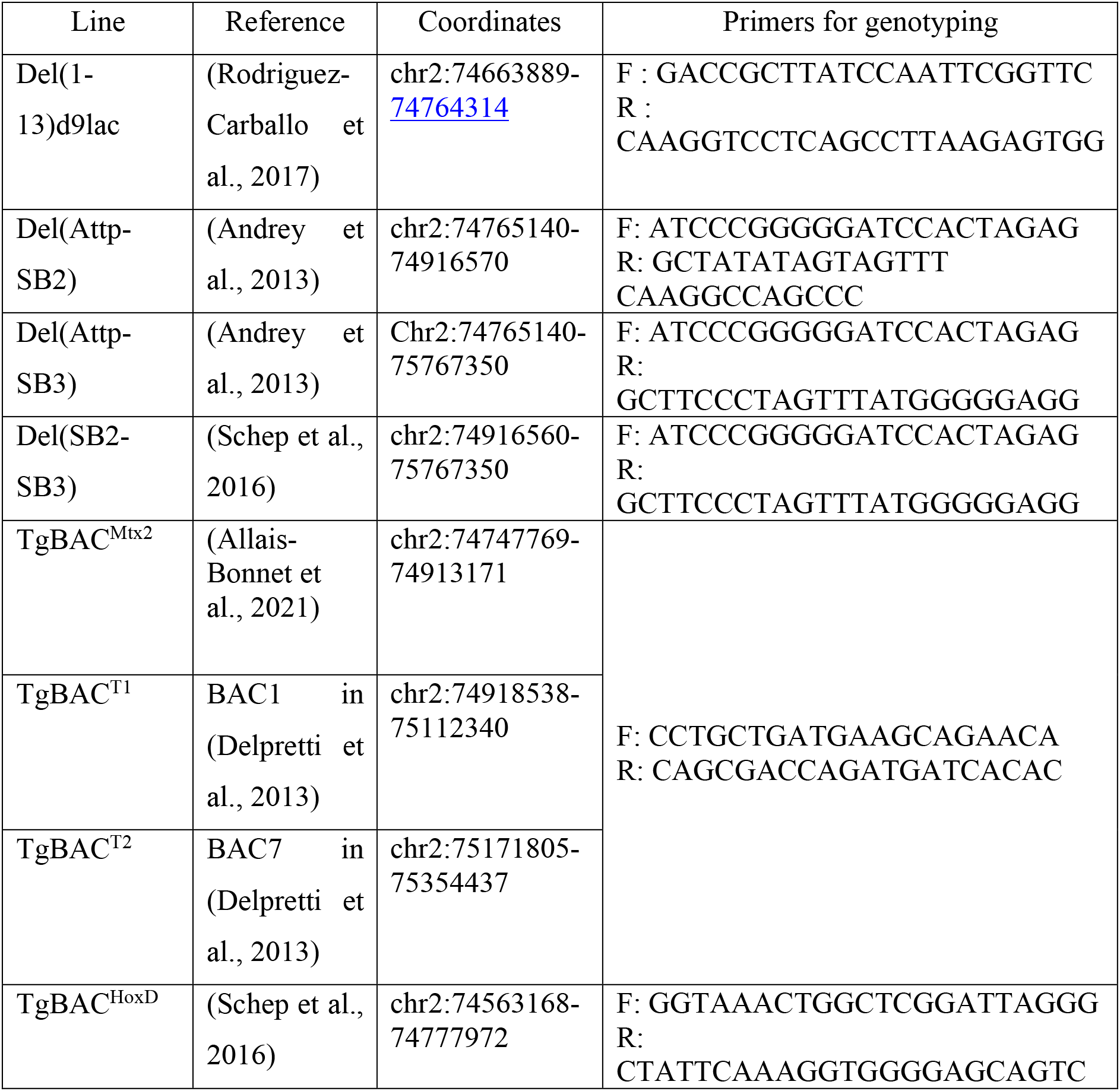
List of pre-existing modified mouse lines.

**Supplementary Table 2.**
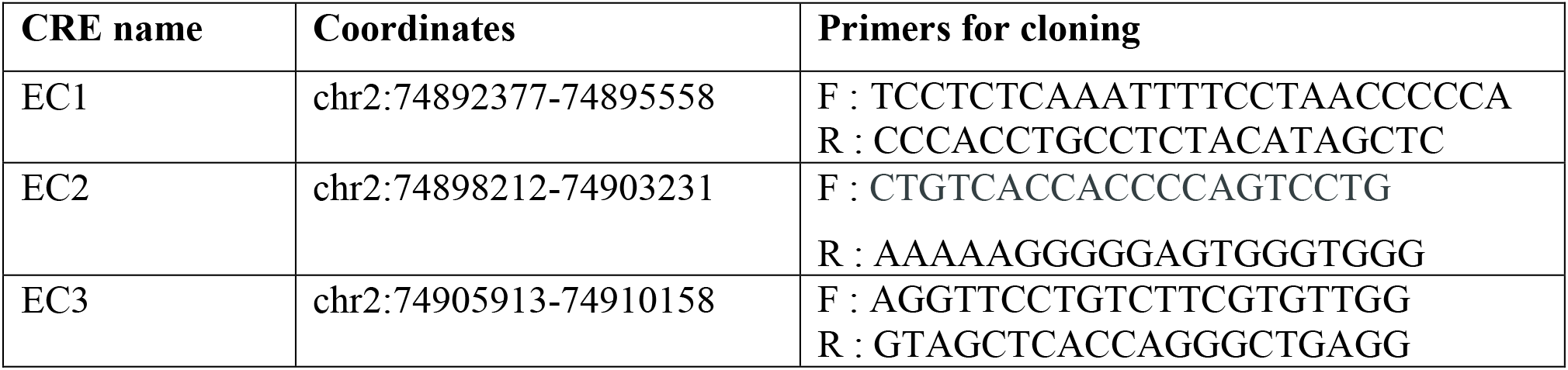
Primers used to clone enhancer candidates.

**Supplementary Table 3.**
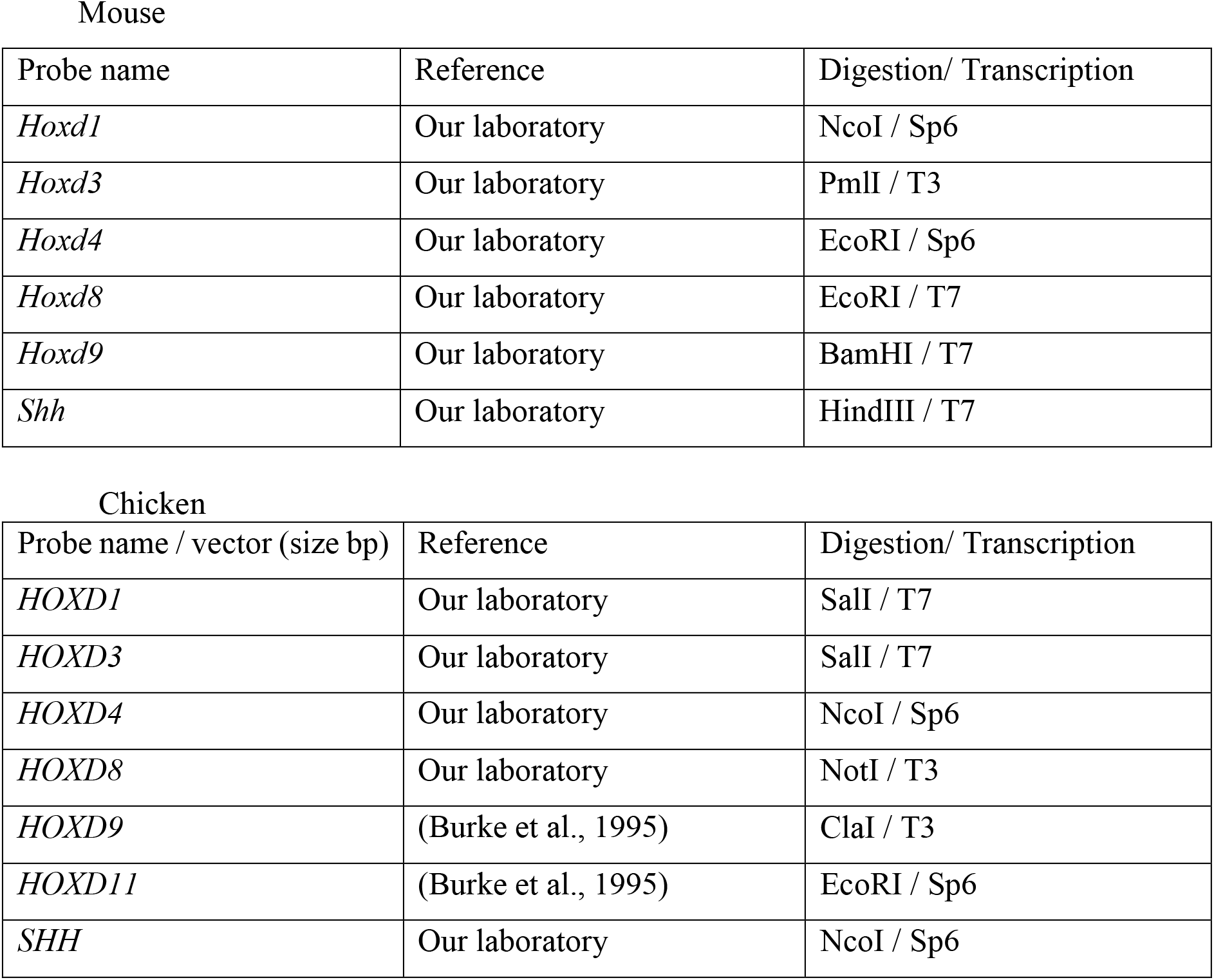
Probes for *In situ hybridization*.

